# ATAClone: Cancer Clone Identification and Copy Number Estimation from Single-cell ATAC-seq

**DOI:** 10.64898/2026.03.11.710984

**Authors:** Lachlan Cain, Anna S Trigos

## Abstract

Single-cell analyses of cancer typically begin by identifying distinct populations of cancer cells by unsupervised clustering. However, in many cases this clustering is explained simply by differences in DNA copy number, which affects the interpretation of differential expression results and tumour heterogeneity studies. To detect and estimate these differences in copy number, we have developed ATAClone.

Applicable to both standalone and multiome scATAC-seq assays, ATAClone first identifies cancer cells with shared DNA copy number profiles (i.e. ‘clones’), then estimates their copy number jointly. Importantly, ATAClone can determine an optimal clustering resolution automatically using simulations. By utilising only stably accessible regions, ATAClone maximises copy number signal while minimising unrelated biological and technical noise. Additionally, by leveraging differences in total DNA between cells, ATAClone can infer absolute copy number, even in the presence of polyploidy.

Using cancer cell mixture experiments, we validate ATAClone’s accuracy in separating clones based on copy number differences. Moreover, using matched scATAC-seq and bulk whole genome sequencing, we show that copy number estimates from ATAClone are more accurate than those derived with existing methods, achieving Pearson correlations between 0.80-0.98 with their bulk-derived estimates and outperforming four other tools that derive copy number from scATAC-seq data. Finally, we show that ATAClone accurately estimates ploidy, including several cases of whole genome doubling which were missed in the matched whole genome sequencing analysis.

ATAClone represents an important tool for disentangling the genetic and non-genetic contributions to gene expression in cancer, providing deeper insight into the evolutionary history and adaptive forces driving a tumour.

## Introduction

Copy number variants (CNVs) are large scale duplications or deletions of a cell’s genome. Here, we refer to cells which share the same set of CNVs as ‘clones’. In cancer, genomic instability and cell stress accelerate the appearance of new clones, while genetic drift and selection (e.g. by drug treatment) shape their relative abundances. Consequently, even a small sample of related cancer cells often consists of multiple clones(1).

With the emergence of single-cell sequencing technologies, fine-grained studies of clones have finally become possible(2,3). However, even when clones are not of direct research interest, such as in studies of transcriptional and epigenetic regulation, clonal copy number differences cannot be ignored. Due to the influence of copy number on gene expression(4,5), DNA accessibility(6), and methylation measurements(7), single-cell analyses of non-genetic phenomena are frequently confounded by pre-existing clonal copy number differences between comparison groups, whether these be unsupervised clusters or experimental treatments. Hence, the identification of clones and their copy number differences is good practice in any single-cell sequencing analysis of cancer.

Several tools have been developed to identify clones and estimate their copy number, both from single-cell DNA sequencing experiments (WGS(8–11) and ATAC-seq(12–16)) and single-cell RNA sequencing experiments(2,4,5,17,18). Typically, these tools obtain copy number estimates for a user-specified set of cells. To identify clones, these estimates are then input into an unsupervised clustering algorithm, the choice of which is also often user-specified. However, the quality of unsupervised clustering, and by extension clonal inference, is tightly linked not only to the quality of the input data, but also its normalisation and the choice of clustering algorithm and its associated hyperparameters. Improved implementations and automation for each of these steps therefore has the potential to increase both the robustness and user-friendliness of clonal inference. Moreover, most existing tools provide an estimate only of ‘relative’ rather than ‘absolute’ copy number and, hence, cannot identify ploidy differences in clones – a more subtle aspect of clonal diversity with implications for evolutionary reconstruction and potential biological significance.

scATAC-seq, as a DNA-based sequencing method, has a more direct relationship between coverage and copy number than scRNA-seq(12), providing better opportunity to derive robust copy number estimates and infer clonal structure from single-cell sequencing experiments. Briefly, ATAC-seq uses Tn5 transposase to selectively fragment and ‘tag’ (‘tagment’) DNA from open chromatin regions for sequencing(19). In the case of 10X Genomics’ scATAC-seq kit, the most popular commercially available implementation, cells are first divided into individual droplets in which the ATAC-seq reaction then occurs separately and is indexed by a unique cell ‘barcode’(20).

Here we describe ATAClone, a tool to infer clonal structure that provides robust ploidy-aware copy number estimates. ATAClone includes automated implementations of quality control, normalisation, clustering, and ‘absolute’ copy number estimation. The power of ATAClone lies in its application of several novel concepts, including: the utilisation of ‘stably-accessible’ regions to measure copy number signal independently of differential accessibility; new metrics for several quality control issues, including unreported coverage biases in the 10X multiome assay associated with cell barcode sequence; an interpretable implementation of graph-based clustering based on Monte Carlo simulation; and the use of total DNA and clone-clone copy number differences for absolute, ploidy-aware copy number fitting. We confirm the robustness and reliability of ATAClone by demonstrating its reproducibility across replicate kidney cancer experiments(21), its sensitivity for clone identification in a labelled lung cancer cell-line mixture experiment(22), and the concordance of its copy-number estimates with those derived from matched bulk WGS in a large cohort of prostate cancer experiments(23). In comparison analyses with similar tools (Copy-scat, epiAneudinfer, RIDDLER, and AtaCNA), we show that ATAClone achieves considerably higher correlation with the ground truth copy number values. Finally, in a comparative analysis with WGS-based copy number estimation tool PURPLE using matched data, we demonstrate that by resolving clonal structure and utilising information on total DNA, ATAClone is able to identify several cases of whole-genome doubling which were missed by PURPLE.

Besides its implications for copy-number calling from scATAC-seq data, our work also provides general advancements in automating several standard single-cell sequencing analyses which are applicable in a range of experimental contexts. All methods described here are collected and implemented in the R package ATAClone, available at https://github.com/TrigosTeam/ATAClone.

## Results

### ATAClone workflow overview

ATAClone infers clonal structure from single-cell ATAC-seq data from cancer samples and provides robust ploidy-aware copy number estimates. ATAClone is designed as a ‘start-to-end’ analysis workflow which provides implementations and guidance for all key analysis steps starting from an unfiltered fragments file from Cell Ranger – users do not need to pre-specify a set of quality cell barcodes for analysis. This contrasts with other copy number estimation tools for single-cell sequencing, where quality control filtering and even the grouping of cells into ‘clones’ is left almost entirely to the user. The structure of the ATAClone workflow consists of steps for **Pre-processing**, **Normalisation**, **Clone Identification**, and **Absolute Copy Number Estimation** (Figure 1).

**Figure 1.**
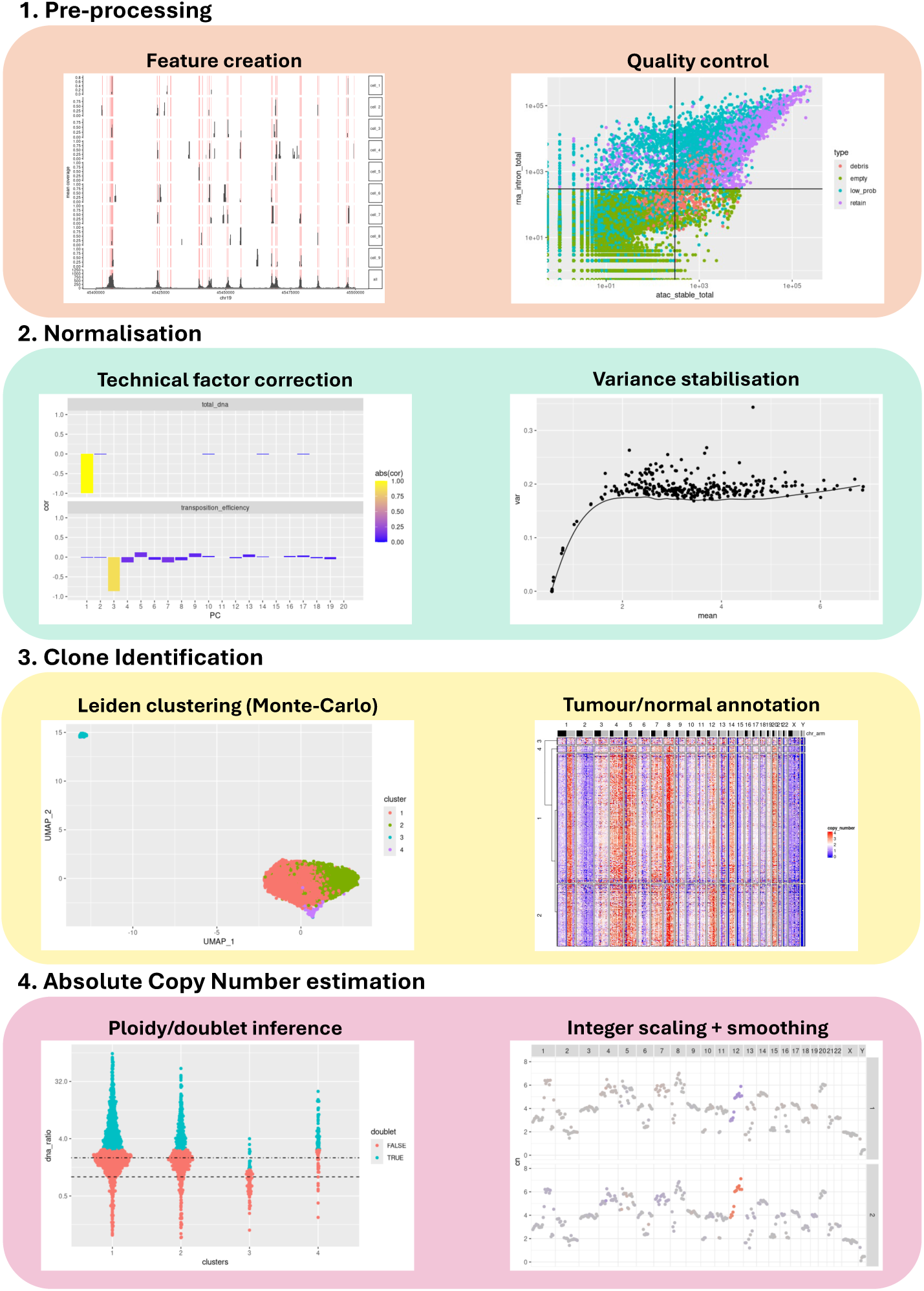
ATAClone workflow overview. The ATAClone workflow consists of four steps. In the first **Pre-processing** step of the workflow, ATAClone performs feature creation by filtering fragments to only those overlapping pre-defined stably-accessible regions, then aggregating these filtered fragments into genomic bins to obtain a bin-by-cell barcode count matrix. The resulting count matrix is subjected to robust **Quality Control** (QC) in which ATAClone filters low-quality cell barcodes based on several QC metrics. In the second step, **Normalisation** is performed. ATAClone first computes statistics to measure and correct technical factors which serve as inputs to a negative binomial regression model. A power transformation is then used for **Variance Stabilisation** of the residuals from the fitted model. These variance-stabilised residuals serve as input to the third step of the workflow, **Clone Identification.** Here, ATAClone automatically identifies clones with shared copy number profiles using a Monte-Carlo approach to optimise Leiden clustering. Comparison of the resulting clusters’ bin count profiles with an external normal reference allows the **Annotation of Tumour and Normal Cells**. The identified normal cells serve as an internal normal reference in the final step of the workflow, **Absolute Copy Number Estimation,** in which differences in total DNA are used for ploidy and doublet inference and copy number ratios (with respect to the internal normal reference) are aggregated at the cluster level to perform integer copy ratio scaling and smoothing. The final ATAClone output is a ploidy-aware copy number estimate for each clone.

### ATAClone analysis is reproducible and robust to sample preparation differences

A challenge of single-cell copy number calling pipelines is the potential for errors in early pre-processing steps to compound through to later analysis steps, leading to false and irreproducible inference of clones and their copy number. To assess this potential sensitivity in ATAClone, we focus initially on the 10X Kidney Cancer replicate datasets. As these datasets use the same tissue but different nuclear isolation protocols (referred to as Chromium Nuclear Isolation, CT sorted, CT unsorted, and SaltyEZ by 10X Genomics), we reason that similar results across replicates would support ATAClone’s robustness and reproducibility, even under different technical conditions.

To test the consistency of ATAClone’s QC filtering statistics to measure and correct the artefacts we hypothesise, including cell debris, variation in transposition efficiency, and variation in sequencing depth, we ran ATAClone across all four replicate experiments under default settings. Across the four nuclear isolation protocols, ATAClone retained similar numbers of cells across the four nuclear isolation protocols (Figure 2a, Supplementary Figure S1a). Notably, the set of cell barcode sequences ATAClone flags as “low probability” demonstrated consistently lower than expected ATAC-seq coverage across all protocols, confirming a predictable coverage bias related to cell barcode sequence in the 10X multiome assay. By contrast, the number of cell barcodes ATAClone flagged as debris varied across protocols; in the CT sorted protocol the artefact was nearly eliminated, as expected given this protocol included an additional FACS step to remove non-cellular material. After normalisation and PCA, ATAClone identified principal components corresponding to the same unwanted technical factors in all four protocols, with total DNA invariably corresponding to PC1 and transposition efficiency corresponding to PC2-PC3 depending on the experiment (Figure 2b, Supplementary Figure S1b). Together, these results support the existence of several recurring artefacts in the 10X multiome assay which ATAClone measures and accounts for.

**Figure 2.**
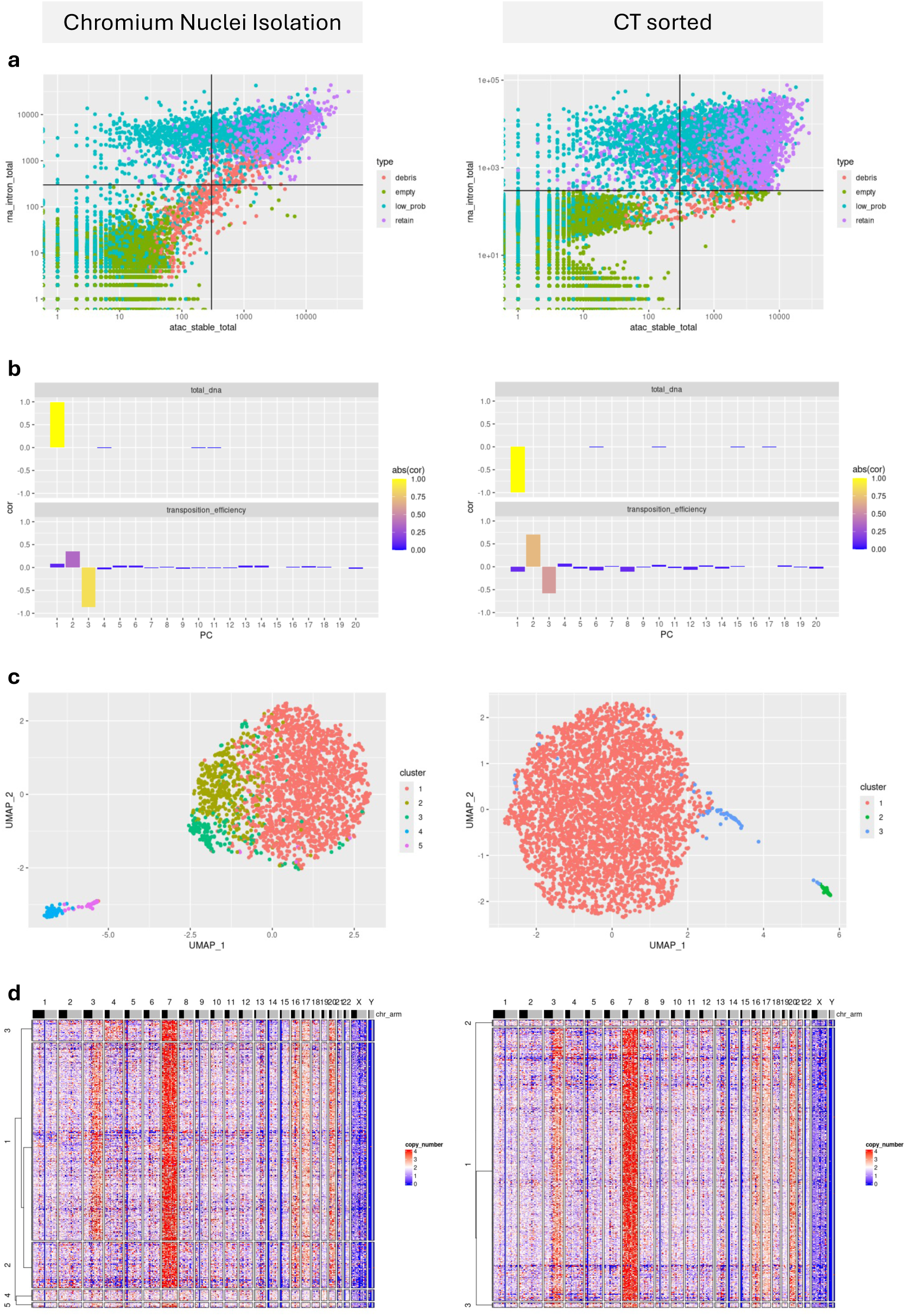
Comparison of ATAClone workflow between 10X Human Kidney Cancer replicate experiments (Chromium Nuclei Isolation vs. CT sorted). Starting from unfiltered data, ATAClone was run to completion for both the Chromium Nuclei Isolation (left column) and the CT sorted (right column) 10X Human Kidney Cancer replicate experiments. **(a)** Using a standardised set of filtering parameters, ATAClone automatically filters cells for several artefacts. Scatter plots show total ATAC-seq stably-accessible fragments (X axis) vs. total RNA-seq intronic UMIs (Y axis) per cell barcode. Horizontal and vertical lines show the minimum X and Y value to be retained for downstream analysis (**300** for both axes in both experiments). Points are coloured by their classification as artefactual (red = nuclear debris, green = empty droplet, cyan = low probability (under-sequenced) cell barcode) or non-artefactual (purple). Note that low probability barcodes (cyan points) represent the same set of cell barcode sequences in all experiments. Additionally, droplets with a fraction of reads in stably-accessible regions less than 0.05 were also filtered (not shown). **(b)** Bar plots showing Pearson correlation of the first 25 principal components after filtering and normalisation with unwanted technical co-variates. The unwanted technical co-variates shown are strongly associated with total DNA (top) and transposition efficiency (bottom), as measured by the total stably accessible ATAC-seq fragments per barcode and the ‘t’ statistic corresponding to transposition efficiency (see: Methods), respectively. Bar height represents the correlation with each co-variate (positive or negative) while bar colour represents its absolute value. **(c)** Scatter plot showing UMAP embeddings (X and Y axes) of filtered barcodes and coloured by unsupervised cluster assignment by ATAClone’s automated clustering approach. Both UMAP embeddings and clusters are computed based on PCA embeddings of the normalised and filtered data after discarding PCs associated with the technical co-variates shown in **(b)**. **(d)** Heatmap showing ‘relative’ copy number estimates (i.e. scaling total DNA to be same as reference cells) across consecutive ∼10MB bins (columns) for each cell barcode (rows), based on an external set of normal reference cells. Cell barcodes are grouped by the same clusters as in **(c)** with clusters sorted by single-linkage hierarchical clustering on the Manhattan distances of their mean bin values. Clusters of non-tumour cells (Clusters 4 and 5 in the Chromium Nuclei Isolation and Cluster 2 in the CT sorted) are distinguishable both as an outgroup in this clustering, and by their relatively constant copy number values. Copy number is clipped to a maximum value of 4 for clearer visualisation.

Reasoning that, as biological replicates, each experiment should at least share the same majority clone, we next sought to characterise and compare the clonal compositions of each replicate experiment. Surprisingly, ATAClone identified differing numbers of clones across replicate experiments, with the Chromium Nuclear Isolation protocol experiment containing 3 clones while the CT unsorted, CT sorted, and SaltyEZ protocol experiments contained just one clone each (Figure 2c, Supplementary Figure S1c). However, visualisation of the copy number ratios at the single-cell level confirmed their homogeneity within each clone as well as the consistency of their clone-clone differences across large contiguous regions, suggesting the detected clones contain genuine copy number differences (Figure 2d, Supplementary Figure S1d). Moreover, ATAClone’s cluster-level absolute copy number estimates displayed clear alignment to integer values, including in regions with clone-clone copy number differences (Supplementary Figure 2). Comparison of these absolute copy number estimates across the replicate experiments additionally supported that each experiment contains the same majority clone, suggesting that the additional clones present in the Chromium Nuclear Isolation protocol are genuine and may arise due to the genetic instability of cancer.

### ATAClone clone identification is highly sensitive

ATAClone’s graph-based clustering algorithm to identify clones aims to automatically identify an optimal resolution parameter for the Leiden algorithm while obtaining type I error rate control via a simulation approach. To confirm that this control is achieved while retaining sensitivity to genuine clonal differences, we use both the scmixology2 lung cancer cell line mixture experiment and the 10X 10k PBMC Multiome Nextgem Chromium Controller dataset as positive and negative controls, respectively.

To assess ATAClone’s sensitivity to known copy number differences, we measured its performance on the scmixology2 dataset which contains labelled mixtures of 5 distinct lung cancer cell lines. Reasoning that, due to genetic instability, some cell lines could contain multiple clones, our performance metrics included two bi-directional measures of clone-label congruity, the Adjusted Rand Index (ARI) and Adjusted Mutual Information (AMI), as well as two uni-directional measures, the homogeneity and the completeness. The latter two measures correspond to the purity of clone assignments to a single cell line label, and the purity of assignment of cell line labels to a single clone, respectively.

Across the five cell lines contained in scmixology2, ATAClone identified 10 tumour clones in total (Figure 3a). These clones achieved a very high homogeneity score with the cell line labels at 0.97, indicating that for each of ATAClone’s clusters nearly all members belonged to the same cell line. By contrast, the completeness score was only modestly high at 0.72 as ATAClone identified multiple clones within some cell lines. As a result, overall clustering metrics Adjusted Rand Index (ARI) and Adjusted Mutual Information (AMI) between ATAClone’s clusters and cell line annotation were only moderately high at 0.71 and 0.72 respectively (Figure 3b). Nevertheless, different clones belonging to the same cell line display stark estimated copy number differences across large contiguous regions, consistent with genuine copy number events (Figure 3c). For example, clusters 5 and 9, both originating from the H2228 cell line, separate by a whole chromosome 18 deletion, while clusters 6 and 8 of the H1975 cell line separate by whole chromosome 4 deletion as well as a gain of the q arm of chromosome 3. Additional unresolved clonal structure is also evident in several clusters, such as cluster 1 (corresponding to the HCC827) cell line where within-cluster sorting reveals whole chromosome gains at chromosomes 4 and 14 as well as a gain of the q arm of chromosome X.

**Figure 3.**
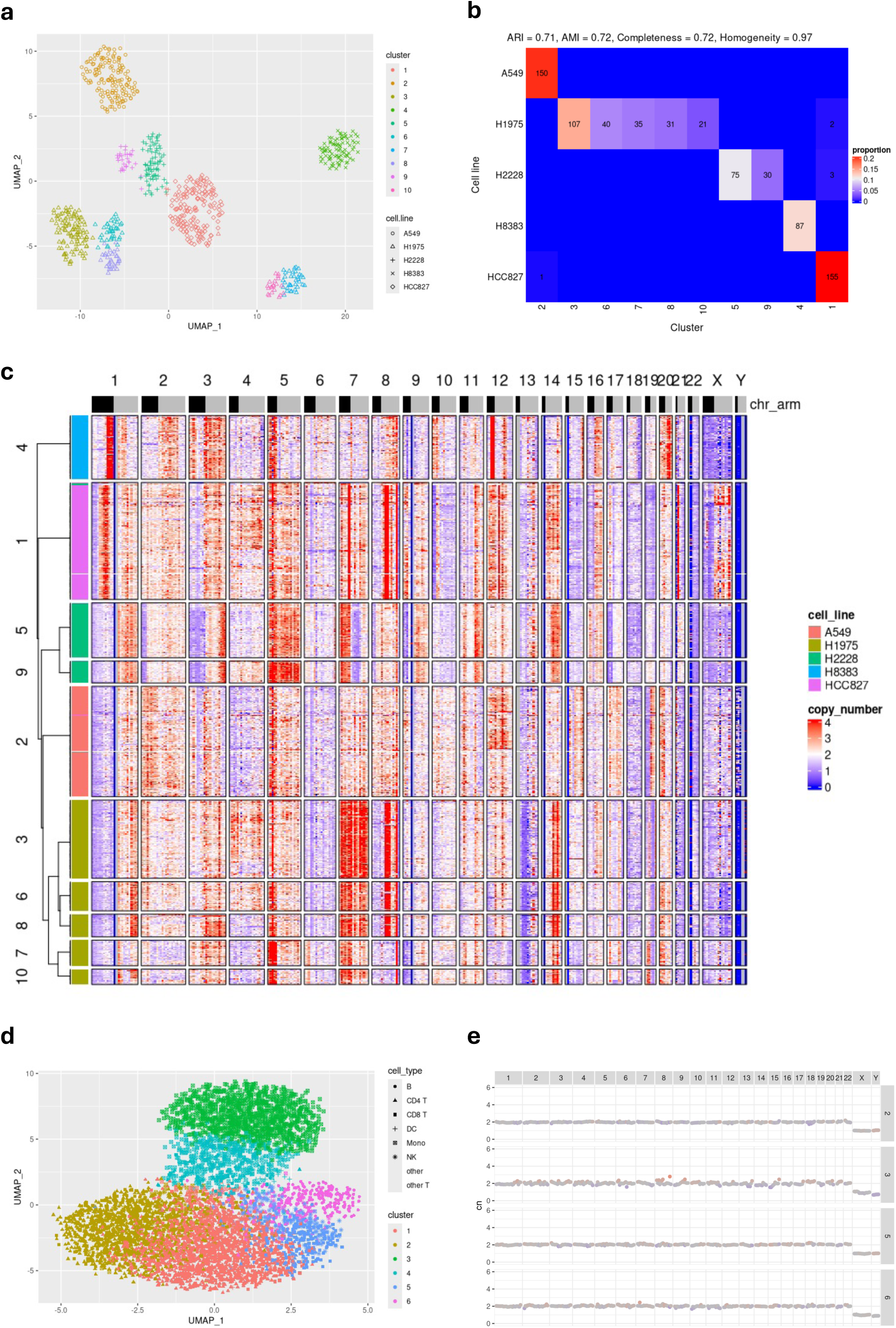
Evaluating specificity and sensitivity of ATAClone’s automated clustering approach. (a-c) To evaluate the sensitivity of ATAClone’s automated clustering approach, ATAClone was run on the scmixology2 lung cancer cell line mixture experiment as a positive control. We used the dataset’s pre-computed cell line labels for each cell barcode based on their SNVs. (a) Scatter plot showing UMAP embeddings (X and Y axes) and coloured by unsupervised cluster assignment by ATAClone’s automated clustering approach. Shapes represent ground truth cell line labels. (b) ATAClone’s clustering demonstrates high purity in the assignment of each of cluster to a single cell type label (homogeneity) but only moderately high purity in the assignment of each cell type label to a single cluster (completeness). Heatmap shows the confusion matrix of unsupervised cluster assignment by ATAClone’s automated clustering approach (X axis) vs ground truth cell line label (Y axis). Each combination is coloured by its overall proportion in the matrix and labelled in text with its frequency (frequencies of zero are omitted). (c) Heatmap showing ‘relative’ copy number estimates (i.e. scaling total DNA to be same as reference cells) across consecutive ∼10MB bins (columns) for each cell barcode (rows), based on an external set of normal reference cells from the human jejunum. Cell barcodes are grouped by the same clusters as in (c) and (d) with clusters sorted by single-linkage hierarchical clustering on the Manhattan distances of their mean bin values. Cell line label is indicated on the left side of the plot. Copy number is clipped to a maximum value of 4 for clearer visualisation. (d-e) To evaluate the specificity of ATAClone’s automated clustering approach, ATAClone was run to on the 10X 10k PBMC Multiome Nextgem Chromium Controller experiment (non-cancer) as a negative control. Under ATAClone’s simulation approach, 95% of barcodes are expected to be assigned to the same cluster in the absence of copy number or differential accessibility signal. (d) Scatter plot shows UMAP embeddings (X and Y axes) and is coloured by unsupervised cluster assignment by ATAClone’s automated clustering approach. ATAClone clusters show strong overlap with the cell type labels. hapes represent cell type predictions by Azimuth based on the scRNA-seq component of the multiome assay. (e) Scatterplots showing ATAClone’s fitted absolute copy number estimates (Y axis) for each consecutive ∼10MB bin along the length of each chromosome (X axis) for the same clusters as identified in (d). Cluster 1 was designated as the reference cluster, and so is not shown. Otherwise, non-consecutive or omitted cluster numbers occur where clusters corresponding to doublets were removed. Points are coloured by their difference in estimated copy number from a reference value for the same bin (here, the bin mean for all clones is the reference), clipped to ±1.5.

Finally, reasoning that non-tumour cells should not generally contain copy number differences, we hypothesised that ATAClone would not identify clones at a rate above the type I error rate in the PBMC experiment, provided it is correctly controlled. Unexpectedly, this was not the case, with ATAClone identifying 6 clusters of non-tumour cells (Figure 3d). Cell typing using Azimuth shows the identified clusters correspond to genuine biological cell types, suggesting this clustering is driven by some residual differential accessibility, as opposed to a lack of statistical control. Nevertheless, copy number estimates for ‘clones’ identified in normal-only samples typically differ only by sub-integer values over non-contiguous regions (Figure 3e), suggesting they are unlikely to confound clonal analysis.

Overall, ATAClone’s automated clustering algorithm demonstrated high sensitivity in identifying tumour clones, as demonstrated by its ability to easily resolve known cell lines, as well as the uncharacterised clonal structure within each cell line.

### ATAClone copy number estimates are highly accurate

We next sought to measure the accuracy of ATAClone’s copy number estimates. For this, we used a cohort of metastatic prostate cancer samples(23), which consists of 24 matched 10X Genomics multiome and WGS data generated from sequential sections of the same tissue block. This dataset contained a total of 7 patients with 2-6 metastatic sites sampled per patient. Copy number estimates were derived from the WGS data using the PURPLE pipeline, as described previously(23). To make the estimates from PURPLE to ATAClone comparable we pseudo-bulked all tumour clones for each ATAClone run after clone-level copy number estimation and assumed a similar clonal composition between each single-cell experiment and its bulk reference. To benchmark ATAClone with other methods, we also ran the scATAC-seq copy number estimation tools AtaCNA(14), Copy-scAT(13), epiAneufinder(12), and RIDDLER(16) for the same set of samples. Additionally, to investigate the impact of restricting analysis to only stably-accessible regions, ATAClone was run in two modes: “stable” in which copy number is estimated only from stably-accessible regions (its default mode) and “all” in which copy number is estimated from all ATAC-seq fragments.

In its “stable” mode, ATAClone’s copy number estimates showed consistently superior performance to other methods (Figure 4A), achieving a median correlation with the ground truth of 0.941, compared to copy-scAT’s 0.861, epiAneufinder’s 0.744, RIDDLER’s 0.738, and AtaCNA’s 0.642. Interestingly, ATAClone’s “stable” mode also generally marginally outperformed its “all” mode despite significantly lower ATAC-seq fragment coverage, achieving a higher correlation value in 19 out of 24 samples. Excluding its “all” mode, ATAClone in “stable” mode also achieved the highest correlation value of all tools in 23 out of 24 samples (Supplementary Figure S3A).

**Figure 4:**
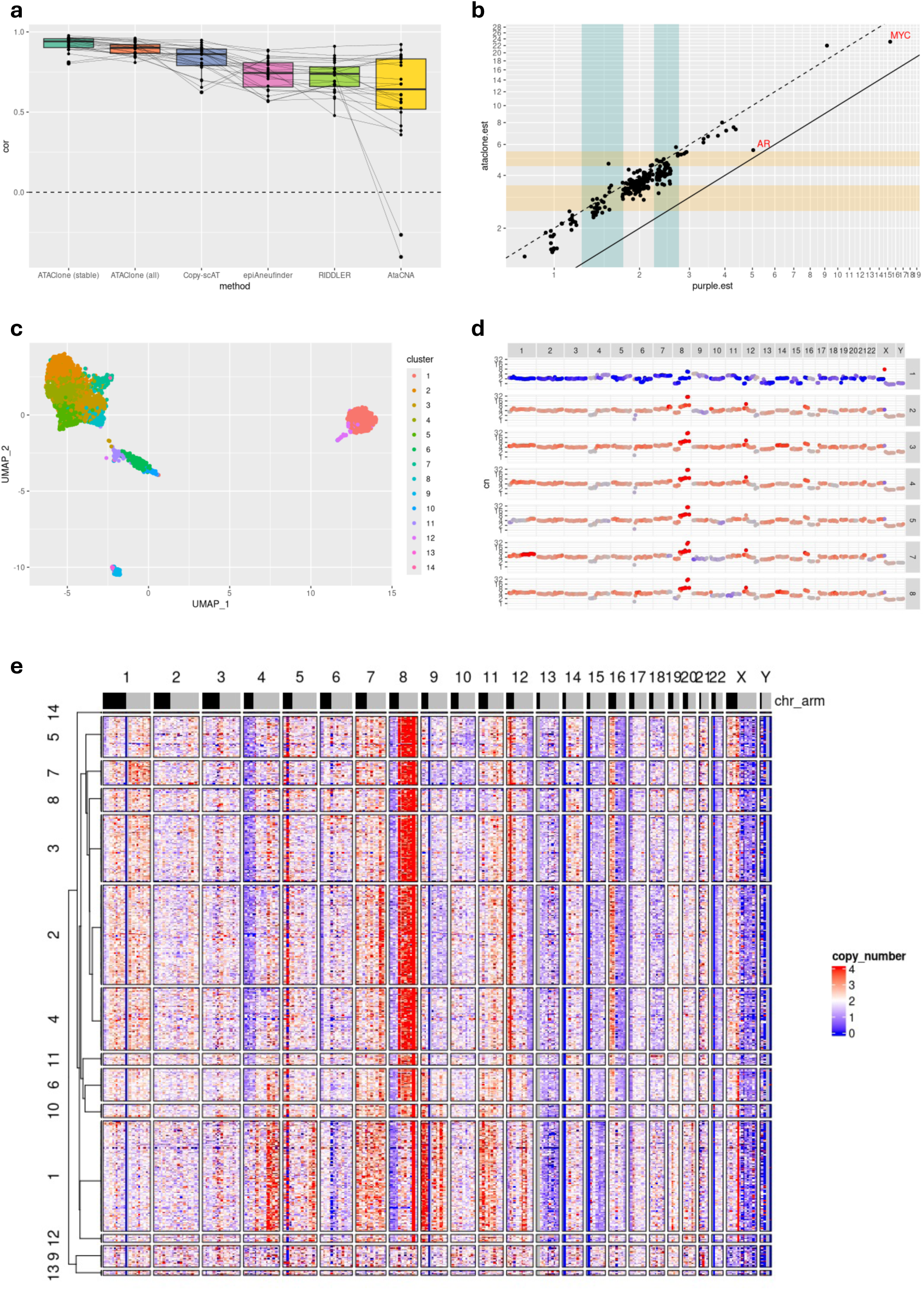
ATAClone accurately detects clonal copy number changes, including in clinically relevant oncogenes. **(a)** Pearson correlation of pseudo-bulked copy number estimates from ATAClone, Copy-scat, epiAneudinfer, RIDDLER, and AtaCNA compared to WGS-derived estimates for matched tissue across 24 metastatic prostate cancer samples. Lines connect the correlation values for the same sample across the boxplots for different methods. Two versions of ATAClone’s copy number estimates are shown: estimates based on fragments at stably-accessible regions only (stable) and estimates based on all fragments (all). **(b-e)** ATAClone results for the CA0034 paraaotic_lymph_node_1 sample. WGS previously identified amplifications in the prostate cancer oncogenes MYC and AR. **(b)** Scatterplot of WGS-derived copy number estimates from PURPLE (x-axis) vs pseudo-bulked single-cell-derived copy number estimates from ATAClone (y-axis), showing strong correlation (cor = 0.954). The bins containing MYC and AR are labelled and correspond to the regions chr8:125150909-135144772 and chrX:61000000-70504090, respectively. Teal vertical bars highlight the inter-integer copy number intervals 1.25-1.75 and 2.25-2.75 (centred on 1.5, and 2.5, respectively) while gold horizontal bars highlight the equivalent doubled intervals 2.5-3.5 and 4.5-5.5 (centred on 3 and 5, respectively). A high density of copy number values in both the teal and gold regions suggests underestimation of ploidy by PURPLE by a factor of 0.5 which is corrected by ATAClone. Parallel lines correspond to y = x (solid), and y = 2x (dashed). **(c)** Scatterplot showing UMAP embeddings (X and Y axes) and coloured by unsupervised cluster assignment by ATAClone’s automated clustering approach. **(d)** Scatterplots showing ATAClone’s fitted absolute copy number estimates (Y axis) for each consecutive ∼10MB bin along the length of each chromosome (X axis) for the same clones as identified in **(c)**. Non-consecutive or omitted cluster numbers occur where clusters corresponding to either non-tumour cells or doublets were removed. Points are coloured by their difference in estimated copy number from a reference value for the same bin (here, the bin mean for all clones is the reference), clipped to ±1.5. The bin containing MYC corresponds to the highest point of copy number gain on chr8 that is seen in all clones, while the bin containing AR corresponds to the highest point of copy number gain on chrX that is seen only in Clone 1. **(e)** Heatmap showing ‘relative’ copy number estimates (i.e. scaling total DNA to be same as reference cells) across consecutive ∼10MB bins (columns) for each cell barcode (rows), based on an external set of normal reference cells. Cell barcodes are grouped by the same clusters as in **(c)** and **(d)** with clusters sorted by single-linkage hierarchical clustering on the Manhattan distances of their mean bin values. Copy number is clipped to a maximum value of 4 for clearer visualisation. Note that, for visual clarity, the x and y axes of **(b)** and the y axis of **(d)** are displayed on a log scale. However, all correlations displayed in **(a)** were computed in the original linear scale.

To investigate the variability in the performance of ATAClone, we specifically compared the ATAClone copy number estimates for the two lowest correlation samples (incidentally from the same patient), the CA0034 liver_left_11 (cor = 0.810) and the CA0034 liver_right_8 (cor = 0.802), with their ground truth values. This analysis revealed a consistent outlier bin at chrX:61000000-70504090 (containing the prostate cancer oncogene AR) which PURPLE detects as highly duplicated but ATAClone reports as copy neutral (Supplementary Figure S3B). However, copy number in other highly duplicated bins, such as chr8:125150909-135144772 (containing the oncogene MYC), was estimated by ATAClone with good accuracy (Supplementary Figure S3B). Hypothesising that the non-uniform distribution of stable regions could induce a coverage bias affecting copy number estimation around AR, we compared these results with ATAClone’s copy number estimates in “all” mode (Supplementary Figure S3C) which showed a notable improvement in reporting this region as duplicated and a higher correlation with the ground truth (liver_left_11 = 0.87, liver_right_8 = 0.85). To qualitatively benchmark detection of copy number changes in oncogenes, we also compared the copy number estimates of AtaCNA, copy-scAT, epiAneufinder, and RIDDLER, to the ground truth for the same regions in the same samples (Supplementary Figures S4A-S4D). While this comparison was not 1:1, due to different bin widths across tools, as well as epiAneufinder filtering the bin containing AR and RIDDLER the entire X chromosome, the results were generally consistent with ATAClone, with the region containing MYC correctly reported as highly amplified and the regions containing (or surrounding) AR frequently underestimated. Interestingly, though reported as diploid by PURPLE, both samples also showed a high proportion of ground truth copy number values centred on 1.5 and 2.5, suggesting that these tumours are instead tetraploid, as reported by ATAClone (Supplementary Figures S3B, S3C).

To illustrate the applicability of ATAClone’s copy number estimates in comparisons of clones, we conducted a full ATAClone analysis of a third sample from patient CA0034: the paraaortic_lymph_node_1. As in the other CA0034 samples (Supplementary Figures S3B, S3C), this sample also exhibited duplications in the regions containing AR and MYC, as well as apparent whole-genome doubling which is not reported by PURPLE (Figure 4B). ATAClone identified a total of 14 clusters in this sample prior to filtering of doublets and non-tumour cells (Figure 4c) with 7 of these identified as tumour clones for absolute copy number calling (Figure 4d). Interestingly, ATAClone reports a large duplication in the bin containing AR only in Clone 1 as well as an additional amplification of chr8q (containing MYC) which is present in all clones except for Clone 1. ATAClone also reports a ploidy difference between Clone 1 (which is approximately diploid) and all other clones (which are approximately tetraploid) (Figure 4d). Heatmap visualisation of the relative copy number estimates at the single-cell level confirms the homogeneity of the copy number profiles of each clone and the consistency of clone-clone copy number differences, including at the bins containing AR and MYC (Figure 4e).

### ATAClone identifies whole-genome doubling

ATAClone’s absolute copy number estimation algorithm aims to produce accurate clone-level integer-centred copy number values, even in the presence of polyploidy. By contrast, other comparable tools aim only to obtain a *relative* copy number estimate. Though we have already found ATAClone’s copy number estimates to be highly *correlated* with their matched ground truth, here we aimed at measuring the consistency of their *scales*.

To measure ploidy inference separately from copy number inference, we define ploidy to be the mean copy number value of a clone. Using the same cohort of 24 metastatic prostate cancer samples, we compare the ploidy reported by PURPLE with the average ploidy reported by ATAClone over all clones. To ensure that copy number values are integer-centred, ATAClone performs an integer-scaling step whereby the simple read-depth ratios for each clone, which are hypothesised to contain ploidy information, are multiplied by a scaling factor. As this scaling step could potentially bias the ploidy estimates, we present both the default ‘scaled’ ploidy estimates as well as the ‘unscaled’ ploidy estimates for reference.

Across the entire cohort of 24 samples, ATAClone’s scaled ploidy estimates were only moderately consistent with PURPLE’s ploidy values with a correlation of 0.42, though this represented an improvement over the unscaled ploidy estimates which showed a correlation of just 0.29 with PURPLE (Figure 5a). Notably, however, ATAClone’s scaled ploidy estimates were highly consistent with the unscaled tumour-normal DNA ratio, whereas PURPLE’s estimates occasionally deviated significantly from these ratios. Moreover, a significant proportion of samples had an ATAClone ploidy estimate which was approximately double PURPLE’s ploidy value. Overwhelmingly, these samples were reported as approximately diploid by PURPLE and originated from the same two patients in the cohort: CA0034 and CA0046. Indeed, these CA0034 samples were previously flagged as exhibiting evidence of undetected whole-genome doubling (Figure 4b, Supplementary Figures S3b, S3c).

**Figure 5:**
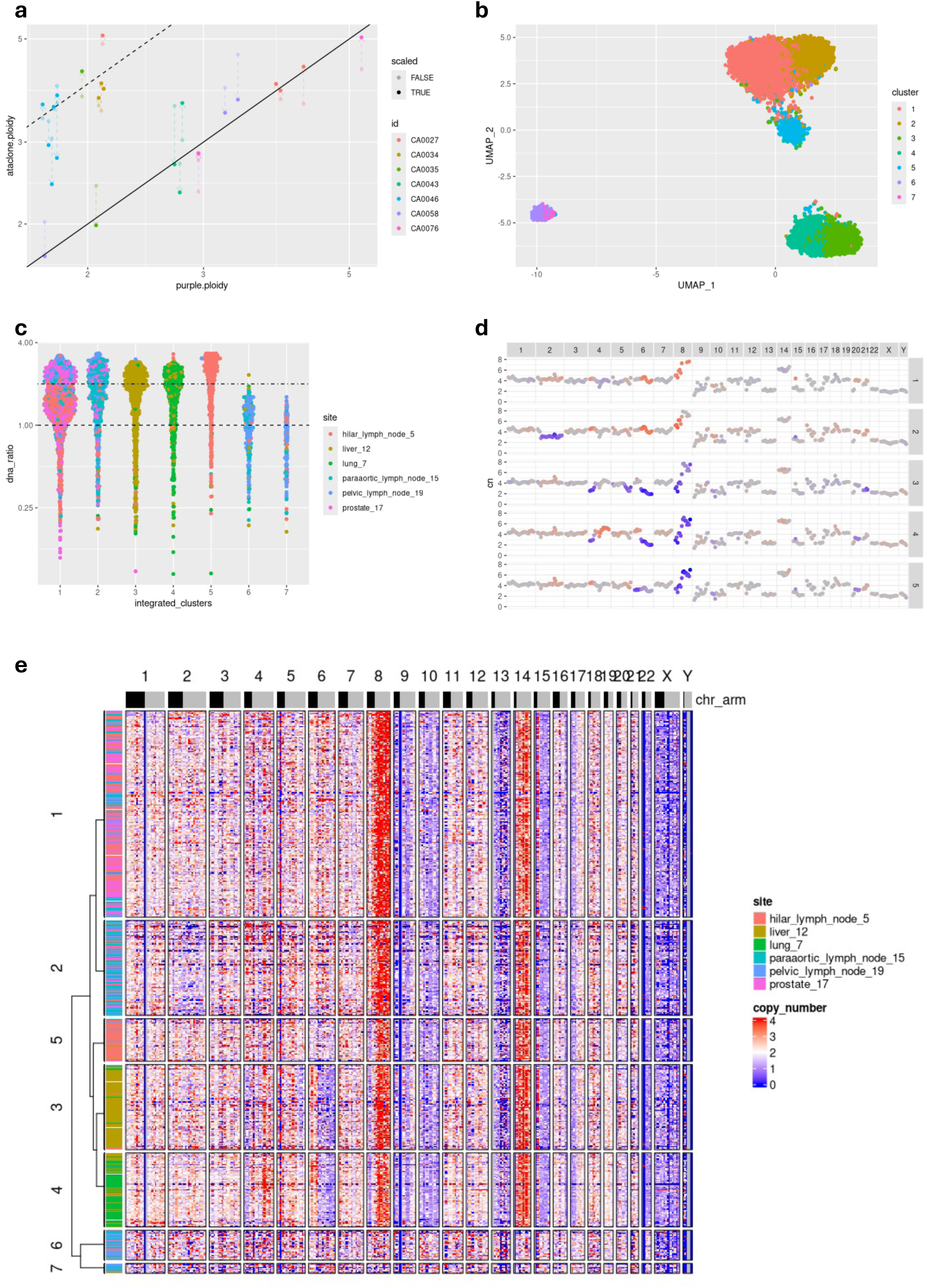
ATAClone can infer polyploidy, even for individual clones. To evaluate the accuracy of ATAClone’s copy number estimates, pseudo-bulked copy number estimates from ATAClone were compared to WGS-derived estimates for matched tissue across 24 metastatic prostate cancer samples with ploidy estimated for each as the mean copy number value. **(a)** Scatterplot shows WGS-derived ploidy estimates from PURPLE (X axis) vs single-cell derived ploidy estimates from ATAClone (Y axis). ATAClone ploidy estimates are generally consistent with PURPLE, though a subset of samples are approximately double in ATAClone. Estimates are coloured by the patient in the cohort they derive from while linked transparent and solid points indicate ATAClone ploidy estimates before and after integer copy number fitting, respectively. Parallel lines correspond to y = x (solid), and y = 2x (dashed). Note that both axes are log-scaled. **(b-d)** To investigate the consistent discrepancy between ATAClone and PURPLE-derived ploidy estimates for patient CA0046, all CA0046 samples were merged and analysed jointly with ATAClone. **(b)** ATAClone successfully integrates multiple CA0046 samples. Scatterplot shows UMAP embeddings (X and Y axes) and coloured by unsupervised cluster assignment by ATAClone’s automated clustering approach. Clusters 6 and 7 were identified as non-tumour cells. **(c)** Beeswarm plot shows the normalised ratio of total ATAC-seq fragments per-cell to the mean total in non-tumour cells (Y axis) for the same clusters as shown in (b) (X axis). Ratios are normalised by scaling for differences in average total ATAC-seq fragments in the non-tumour cells of each sample. All CA0046 tumour clones have approximately double the DNA of non-tumour cells from the same samples. Ratios are coloured by the sample they originate from. **(d)** Clone-level absolute copy number estimates show evidence of whole-genome doubling. Scatterplots showing ATAClone’s fitted absolute copy number estimates (Y axis) for each consecutive ∼10MB bin along the length of each chromosome (X axis) for the same clones as identified in (b). Large contiguous regions of odd-numbered copy number values suggest a failure in ploidy estimation by PURPLE which is rectified by ATAClone. Non-consecutive or omitted cluster numbers occur where clusters corresponding to non-tumour cells were removed. Points are coloured by their difference in estimated copy number from a reference value for the same bin (here, the bin mean for all clones is the reference), clipped to ±1.5. **(e)** Odd-numbered copy number regions are not an artefact of unresolved clonal mixtures. Heatmap shows ‘relative’ copy number estimates (i.e. scaling total DNA to be same as reference cells) across consecutive ∼10MB bins (columns) for each cell barcode (rows), based on an external set of normal reference cells from the human jejunum. Cell barcodes are grouped by the same clusters as in (b-d) with clusters sorted by single-linkage hierarchical clustering on the Manhattan distances of their mean bin values. Sample label is indicated on the left side of the plot. Copy number is clipped to a maximum value of 4 for clearer visualisation.

To further investigate this discrepancy in ploidy between ATAClone and PURPLE, we elected to focus on the samples originating from patient CA0046, representing the majority of outlier samples. After initial pre-processing by ATAClone, including removal of doublets, we merged the bin count matrices of all CA0046 samples to integrate and re-analyse them jointly with ATAClone. Ignoring non-tumour cells, ATAClone identified a total of 5 clones across the six CA0046 samples. Consistently, the total DNA of tumour clones (clusters 1-5) were approximately double that of non-tumour cells (clusters 6-7) after adjusting for differences in sequencing depth between sites (Figure 5c). Interestingly, Clone 1 appeared to exhibit a bimodal distribution centred around a DNA ratio of 1 and 2, suggesting a potential mixture of diploid and tetraploid cells. This evidence of polyploidy was further supported by ATAClone’s absolute copy number estimates with several clones exhibiting large contiguous regions of copy number 3, including chromosome 2 in Clone 2, chromosome 21 in Clone 3, and chromosome 6 in Clone 5 (Figure 5d). As copy number 3 could also occur due to subclonal mixtures of copy number values, we visualised the single-cell-level relative copy number estimates across all clones but found these regions were consistently homogeneous within clones with no evidence of unresolved clonal structure (Figure 5e). Together, these results indicate that ATAClone can detect polyploidy, even at the level of individual clones, at a greater accuracy than that obtained by standard WGS pipelines.

## Discussion

In this work we introduce ATAClone, a highly automated pipeline for the identification and estimation of clonal copy number variants for single-cell ATAC-seq. By reducing the user input required for quality control, clustering, and copy number estimation, ATAClone both streamlines and significantly improves the robustness of a critical analysis in single-cell cancer studies. We demonstrated the reproducibility of the ATAClone analysis workflow, its sensitivity in identifying groups of cells with clonal copy number variants, and the accuracy of its copy number estimates. While ATAClone achieved more accurate copy number estimates than comparable tools for the same group of cells, it also significantly improved on their utility by automatically identifying clonally related cells and their absolute copy number – features not implemented in any comparable tool.

Interestingly, ATAClone also demonstrated superior detection of whole-genome doubling compared to the PURPLE pipeline on matched WGS. This was supported by multiple lines of evidence, including a corresponding increase in the tumour-normal DNA ratio as measured by scATAC-seq, clear alignment of ATAClone’s clone-level copy number estimates to odd numbers over large contiguous regions, and, in some cases, even the PURPLE results themselves when a large proportion of non-integer copy number values was evident. This was surprising given the strong performance of PURPLE for ploidy inference in independent benchmarking(24). However, this same benchmarking also demonstrated a major reduction in performance when whole genome doubling had only recently occurred. That finding appears consistent with our results, where two patients recurrently identified as having whole genome doubling by ATAClone but not PURPLE also showed evidence of mixtures of tumour ploidies across clones, suggesting this doubling had only recently occurred. The detection of clonal ploidy differences with ATAClone are particularly valuable given the known role of tumour ploidy in drug resistance(25) and further enables cancer genomics studies into clonal evolution.

While identifying clones and their copy number differences is good quality control practice in any single-cell analysis of cancer, we envision that ATAClone will also enable more specialised studies of clonal biology. These may include the interaction of copy number changes with transcriptional programs or epigenetic remodelling, or the occurrence of polyploidy in response to drug treatment and the transcriptional programs which arise to support it. The work described here is also significant for its general advancements in automating single-cell sequencing analysis, which is critical as the size of single-cell studies continues to exponentially increase. Notably, we introduced an approach to automatically select an appropriate resolution parameter for graph-based clustering – a finding with significant implications given the ubiquity of graph-based clustering in single-cell analyses(26–28). We also introduce the concept of stably-accessible regions, an ATAC-seq counterpart to stably-expressed genes(29), and demonstrate their utility for copy number estimation. Though bins computed from stably-accessible regions showed some sensitivity in identifying non-copy number-related differences in non-cancer samples – suggesting they are not truly stably-accessible – this did not appear to be produce false positives in the cancer setting where clone identification was found to be highly sensitive. Additional refinement of the criteria for identifying stably-accessible regions may improve their performance in this regard. Finally, we also provide new technical insights into the scATACseq assay, where we uncovered a novel systematic bias in the 10X multiome assay wherein particular cell barcode sequences have reproducibly lower ATAC-seq coverage.

In summary, ATAClone is an R tool that identifies clones and estimates ploidy-aware copy-number. Beyond improvements in performance, ATAClone is set apart from other similar tools in its ability to derive ploidy, even at the level of individual clones, enabling calling absolute copy-number estimation, and its significant automation, which enables robust sample pre-processing and derivation of clonal structure without ad hoc decisions by the user.

## Materials and Methods

### ATAClone method

The ATAClone workflow can be divided into four stages: **Pre-processing**, **Normalisation**, **Clone Identification**, and **Absolute Copy Number Estimation**.

#### 1. Pre-processing

##### Identifying stably-accessible regions

Instead of performing peak calling to identify accessible DNA regions for each sample, ATAClone uses a pre-computed list of regions which are stably-accessible across many different cell types. The rationale of this approach is to minimise non-genetic signals, such as differential accessibility, for better copy number inference.

Here, stably-accessible regions were defined from a study which derived consensus peaks from DNA-seq, and subsequently tested the presence-absence of different peaks across many different samples from 16 organ systems using the Altius DNAse I hypersensitive site index(30), resulting in 76,951 stably-accessible peaks. In ATAClone, fragments were considered to overlap these regions only when the fragment start, end, or both start and end intersect at least one stably accessible region, as proposed in the paired-insertion count method(31). Note that the unfiltered version of this peak set is also used by the scATAC-seq copy number calling tool RIDDLER(16).

##### Binning of ATAC-seq fragments

To improve the copy number signal-noise ratio of features, ATAClone aggregates ATAC-seq fragments into genomic bins to obtain a bin by cell barcode count matrix. For quality-control purposes, ATAClone computes two count matrices: one derived only from fragments overlapping stably-accessible regions and one derived from all fragments.

As a fixed bin width cannot perfectly divide the length of all chromosome arms, resulting in large bin width inequalities, ATAClone instead sets a ‘target’ bin width (default 10Mb). In this approach, bins are obtained by dividing each chromosome arm into n_i_ equiwidth bins, where n_i_ is the number of bins which minimises the absolute difference from the target bin width for chromosome arm i. Fragments are then counted under a bin only when the fragment start site falls within the bin. If a fragment has ‘PCR duplicates’, as inferred by sharing a start and end site, then these duplicates are discarded.

Except where stated otherwise, only the stably-accessible region count matrix is used for downstream tasks such as clustering and copy number estimation.

##### Quality Control

To aid downstream cell filtering and normalisation, ATAClone computes several quality control statistics. These statistics are designed to be intuitive, discriminating, and fast to compute.

While thresholds for all QC metrics can be set manually, ATAClone also implements default values for each. For all default values, and any dataset-specific deviations used in this Article, refer to “Data Pre-Processing” in the Methods.

##### Identifying Empty Droplets

To aid discrimination of nucleus-containing droplets from “empty” droplets(32), ATAClone uses the total fragments in stably-accessible regions per droplet as a proxy for nuclear DNA content. In the case of 10X scRNA-seq + scATAC-seq multiome data, ATAClone additionally computes the total intronic RNA UMIs per droplet as a proxy for nuclear RNA content, as previously described(23,33). The total RNA intronic UMIs is also useful for identifying ‘low-RNA, high-DNA’ droplets, a potential artefact.

##### Measuring transposition efficiency

To measure variation in transposition efficiency across droplets, ATAClone computes the fraction of fragments in stably-accessible regions (as a proxy for fraction of reads in peaks) per droplet. Similar statistics have previously been shown to correlate with variation in the transposase concentration(34) and exposure time(35) in bulk ATAC-seq.

In some cases, the fraction of fragments in stably-accessible regions is insufficient to measure subtle differences in transposition efficiency between cell barcodes. Therefore, ATAClone takes an additional Poisson regression approach to measure transposition efficiency as follows. For each cell barcode_j_, a Poisson regression of the form

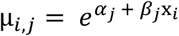

is fitted, where μ_i_ is the expected count for bin_i_ in cell barcode_j_ and x_i_ is the expected count of the same bin in an external reference set of non-tumour cells. This relationship is monotonically increasing in the absence of copy number variants. We note that, in the absence of copy number changes, the expected value of a bin is strongly correlated with fraction of reads in stably-accessible regions for that bin. Therefore, the resulting fitted values, α_j_ and β_j_, correspond to total DNA and the strength of dependence of bin coverage on fraction of fragments in a bin, respectively. Lower β_j_ values correspond to slower growth in coverage with respect to its expected value (i.e. lower transposition efficiency), while higher β_j_ values correspond to faster growth (i.e. higher transposition efficiency). For clarity, we will denote the transposition efficiency as estimated by β_j_ as *t*.

##### Identifying cellular debris

To measure random loss of whole chromosomes due to cell death, ATAClone computes the ‘excess’ zero bin counts per droplet as the observed number of zero bin counts minus the expected number of zero bin counts under a model of ‘cellular homogeneity’ – that is, where the expected proportion of each bin does not vary between cell barcodes.

##### Identifying “low coverage” cell barcodes

10X scRNA-seq + scATAC-seq multiome data exhibits a strong discordance between total RNA UMIs and total ATAC-seq fragments per-cell barcode). Our own analyses strongly suggest that this is an artefact resulting from certain cell barcode sequences which are systematically biased to have lower than expected ATAC-seq coverage.

Reasoning that, due to their lower coverage, these cell barcodes will also have zero detectable ATAC-seq fragments more often, ATAClone measures this bias indirectly by estimating a ‘barcode probability’ – the probability that a cell barcode will have one or more detected ATAC-seq fragments in any given 10X scRNA-seq + scATAC-seq multiome assay. To aid interpretability, ATAClone normalises the estimated barcode probability by multiplying by 735,320 – the total number of possible barcodes in the multiome assay. Therefore, the expected barcode probability if all cell barcodes are equally likely to appear is 1 / 735,320. Hence, the resulting normalised barcode probability estimate is referred to as a ‘barcode odds’.

Given a list of barcode sequences with at least one ATAC-seq fragment from one or more experiments and a complete barcode whitelist, ATAClone estimates a barcode probability as follows. First, each 16 base cell barcode sequence is decomposed into three subsequences: a left subsequence (bases 1-7), a linker subsequence (bases 8-9), and a right subsequence (bases 10-16). This was based on previous findings relating barcode subsequences to low coverage(36) and a related fix pushed by 10X (https://github.com/10XGenomics/cellranger-atac/blob/main/mro/atac/stages/processing/cell_calling/remove_barcode_multiplets/_init_.py).

Based on the Cell Ranger ARC barcode whitelist, these subsequences have 1920, 4, and 1534 possible variations, respectively. However, each of the four linker subsequences associates exclusively with a non-overlapping subset of left and right subsequences. Therefore, a left and right subsequence may only ligate if they associate with the same linker subsequence. For each left subsequence, ATAClone then estimates a left probability P_L_ as

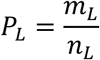

where m_L_ is the frequency of the left barcode and n_L_ is the sum of the frequencies of all left barcodes. A right probability, conditional on the linker subsequence, is also computed as:

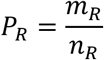

where m_R_ is the frequency of the right barcode and n_R_ is the frequency of all right barcodes with the same linker subsequence. The final barcode probability is then computed as

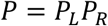

#### 2. Normalisation

##### Technical factor correction and variance stabilisation

ATAClone assumes bin counts to follow a Gamma-Poisson distribution NB(μ, α) – a common model in the bioinformatics literature for a range of sequencing analyses. Across different features, Gamma-Poisson variance is heteroscedastic with variance

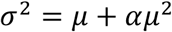

Removing this heteroscedasticity has been shown to improve performance of unsupervised clustering(37).

To account for unwanted technical factors, ATAClone uses the *glmGamPoi* R package to fit the negative binomial regression model

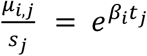

for each bin_i_ where t_j_ is the transposition efficiency for cell_j_ as previously estimated and s_j_ the size factor for cell_j_ as estimated from its total ATAC-seq fragments which is treated as an offset. For model fitting, overdispersion α is set to a fixed value of 0.01.

To correct for differences in transposition efficiency and size factors while stabilising the variance of bin counts, ATAClone computes a corrected and normalised count z from x and the fitted regression model as

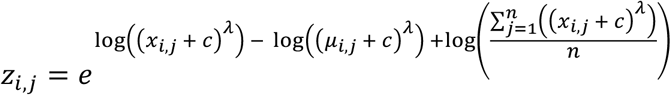

where μ_i,j_ is the fitted mean for bin_i_ of cell_j_ and λ and c are a user-specified fractional power and pseudo-count, respectively. Briefly, the transformation (*x_i,j_* + *c*)*^λ^* serves to stabilise the variance of each bin_i_, the use of a log-scale when subtracting (*μ_i,j_* + *c*)*^λ^* generates residuals corrected for differences in s_j_ while preserving the ratio of x_i,j_ to μ_i,j_, and the final term acts to centre these residuals on each bin’s overall mean value before transforming back to the original scale. By default, ATAClone uses λ = 0.4 and c = 0.5. To assess the fit of the gamma-Poisson model, ATAClone simulates a corresponding ‘null’ – that is, without biological variation – matrix of counts y_i,j_ where y_i,j_ ∼ NB(μ_i,j_, α) and applies the same normalisation procedure, after which the mean-variance relationships of the simulated and original data are compared to confirm a fit.

##### KNN graph assembly

To denoise the data, ATAClone then performs dimensionality reduction using the *irlba* implementation of sparse PCA(38)with all trailing components after PC20 discarded.

Finally, based on Euclidean distance in their PCA embeddings, ATAClone builds a KNN graph of cell barcodes (here, k = 20) for both the simulated and unsimulated which serves as the input to the graph-based clustering procedure.

##### Graph-based clustering

Using the previously described normalisation and KNN graph assembly procedure, ATAClone obtains KNN graphs from both the simulated and unsimulated count matrices. These graphs serve as inputs to the *igraph*(*39*) implementation of the Leiden clustering algorithm(40) with the constant Potts model (CPM) resolution parameter(41).

To automatically identify an optimal CPM resolution parameter for the unsimulated data, ATAClone takes a probabilistic approach using the simulated data as follows. Let γ_1_, γ_2,_ γ_3_… γ_k_ be an ascending sequence of resolution parameters and p_1_, p_2,_ p_3_… p_k_ be the probabilities at each resolution that a single graph node will **not** be assigned to the largest cluster by the Leiden algorithm under the null model – that is, in the simulated data. Given a maximum acceptable type I error rate α (here, 0.05), the optimum resolution can be found as the maximum γ_i_ for which p_i_ < α. As the Leiden algorithm is stochastic, for each resolution γ_i_, ATAClone estimates p_i_ from the simulated data over n_iter_ independent Leiden clustering iterations (here n_iter_ = 500,000 / n_nodes_).

Once an optimum resolution is found on the simulated data, the algorithm stops and this resolution is applied to the unsimulated data to find a single initial Leiden clustering. The algorithm is then applied recursively with three modifications:

1. in the count matrix simulation step, bin proportions estimates p_j_ are updated and computed separately based on each of the previously identified clusters in the unsimulated data

2. in the normalisation and graph-assembly step, to keep cell-cell distance in the unsimulated data constant across recursion steps, PCA is performed using all members of all clusters as input, but KNN graph assembly is separate for each of the previously identified clusters in both the simulated and unsimulated data

3. in the graph-based clustering step, separate optimum resolutions are found for the KNN graphs of each of the previously identified clusters

In the results shown in this study, just one recursive step is used after the initial clustering for performance reasons.

#### 4. Absolute copy number estimation

##### Internal reference annotation

For absolute copy number estimation, ATAClone requires non-tumour cell barcodes in the same experiment to use as an ‘internal reference’. To identify these cell barcodes, an external reference sample of known non-tumour cells is used to obtain a relative copy number estimate of each bin in each cell barcode. This is computed as the proportion of counts in a bin divided by the mean proportion of that bin in the reference, multiplied by 2 for all autosomes (or all chromosomes if the reference is female). The internal reference is then selected as one or more of the previously identified clusters which exhibit a flat relative copy number.

##### Doublet detection

To ensure that copy number and ploidy inference is not confounded by doublets, scDblFinder(42) was run using a custom transformation of (*x* + *c*)*^λ^* (i.e. variance stabilisation without library size correction) to identify doublets, with no pre-computed clusters given and clustering disabled. As scDblFinder’s “score” is interpretable as a binary classification probability, scores greater than 0.45 were considered as doublets to be slightly more conservative than the typical threshold of 0.5.

##### Integer copy number scaling

Absolute copy number is then estimated for each clone in two stages. In both stages, the same range of “scaling factors” s_1_, s_2_,… s_k_ are considered.

In the first stage, an initial copy number estimate x_j_ is obtained for each bin_j_ by dividing its mean by the mean of in internal reference, multiplied by 2 for all diploid chromosomes in the reference. For each bin_j_, an estimate of the variance of the means is obtained by bootstrapping, and the inverse of this bootstrapped variance is used as a precision weight w_j_. The initial copy number estimates and their precision weights are then input into the Ckmeans.1d.dp algorithm to identify preliminary clusters of copy number levels. A second set of weights v_j_ are then computed as the inverse of the frequency of each bin’s preliminary cluster (i.e. rarer clusters are given more weight). Then, for each scaling factor s_i_ a weighted sum of squared errors is computed as

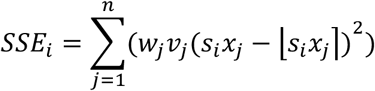

where ⌊⌉ denotes an integer rounding operation. If just one clone is present, then the scaling factor with the lowest sum of squared errors is used. The resulting absolute copy number estimate is then the preliminary estimate multiplied by this scaling factor. Otherwise, we consider all possible combinations of tumour clones. Let the preliminary copy number estimate of bin_j_ be called x_j_ in the first clone considered and y_j_ in the second. For each combination, we compute the ratio of x_j_ / y_j_ for all bins, then use bootstrapping of this ratio to obtain a new precision weight w_j_ for each bin_j_. These ratios and their weights are then input into the Ckmeans.1d.dp algorithm to identify preliminary clusters of pairwise copy number differences. A second set of weights v_j_ are then computed as the inverse of the frequency of each bin’s preliminary cluster. Then, for each combination of scaling factors sf_1_ and sf_2_ corresponding to the two clones being compared, a weighted sum of squared errors is computed as

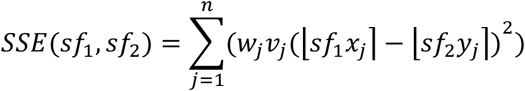

This is repeated for all possible sf_1_ and sf_2_ values, iterating over s_1_, s_2_,… s_k_. For each scaling factor of each clone, its final sum of squared errors is its error from the first stage of the algorithm (weighted by the frequency of members of its clone) plus the sum of its errors in the second stage of the algorithm across all pairwise comparisons involving that clone (weighted by the minimum frequency of the two clones compared). For each clone, the scaling factor with the lowest sum of squared errors is used and its resulting absolute copy number estimate is then the preliminary estimate multiplied by this scaling factor.

##### Copy number smoothing

After integer copy number scaling, copy number values are smoothed as follows. For each bin_i_ a vector of distance weights 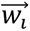 is computed as

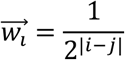

where |i – j| is the absolute difference in indices between two bins. Note that weights are set to 0 for any i and j not corresponding to bins on the same chromosome.

Additionally, for each bin_i_ a vector of outlier weights 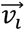 is computed as

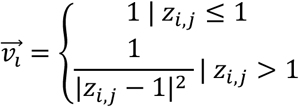

where z_i,j_ is a z score of the log-odds ratio of the clone-level copy ratio at bin_i_ compared to bin_j_. A smoothed copy number ratio is then estimated as

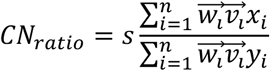

where s is the fitted scaling factor for a clone, x_i_ is the mean count for bin_i_ in a clone and y_i_ is the mean count for bin_i_ in the reference cells. This is multiplied by 2 for all chromosomes which are diploid in the reference to obtain a smoothed copy number estimate.

##### Data pre-processing

As input, ATAClone uses the fragments file output by Cell Ranger ATAC or Cell Ranger ARC.

##### Cell barcode filtering

For both the standalone 10X scATAC-seq and 10X scRNA-seq + scATAC-seq multiome assays, the following filtering steps were applied. To remove “empty droplets”, cell barcodes with 300 or fewer fragments were removed. To reduce the impact of differences in transposition efficiency on copy number inference, cell barcodes with a fraction of reads in stably-accessible regions less than 0.05 were also removed. Finally, to reduce false copy number signal from random loss of whole chromosomes, cell barcodes with 13 or more ‘excess’ zero bin counts were removed. Each threshold was chosen based on generalisable good performance across all samples tested.

In the case of 10X scRNA-seq + scATAC-seq multiome data, the following additional filtering steps were applied. Firstly, to remove “low-RNA high-DNA” cell barcodes, and to further refine the removal of “empty” droplets, cell barcodes were filtered for a minimum total RNA intronic UMIs of 300. Secondly, to reduce the influence of technical variation in ATAC-seq coverage on clonal inference and ploidy estimation, cell barcodes with a ‘barcode odds’ of 1 or less were removed. We note that, because the barcode odds are pre-computed, the same cell barcode sequences were removed in all samples based on this criterion.

In the case of standalone 10X scATAC-seq data only, cell barcodes flagged as ‘barcode multiplets’(36) were removed based on the ‘singlecell.csv’ file output by Cell Ranger ATAC (v).

##### Other Pre-processing Parameters

In this study, during normalisation the overdispersion parameter α was set to a 0.01 for all samples except for the 10X 10k PBMC Multiome Nextgem Chromium Controller dataset where it was set to 0. For a single metastatic prostate cancer sample (CA0027 paraaortic_lymph_node_1), Leiden clustering was not performed recursively to reduce number of clusters for copy number fitting. During clustering, the seed for Leiden clustering was altered for a minority of samples to better separate clusters corresponding to doublets. During doublet filtering, the total DNA threshold for doublet filtering was overridden in some clusters in a minority of samples to obtain a better boundary between genuine cells and apparent doublets.

##### Comparison of scATAC-seq copy number tools

All tools were run in their default configurations and according to their documented methodology where possible. However, substantial modifications were made to AtaCNA code so that it would run to completion including disabling GC correction which was found to prevent run failure during segmentation. EpiAneufinder was also modified from its default configuration to set the minimum number of fragments to 10,000 (as the default 20,000 resulted in removal of most cells in most samples) and increasing the bin width to 1MB based on advice in the tool documentation for lower coverage samples.

##### Comparison with PURPLE

PURPLE was run for each WGS sample as previously described(23). As PURPLE outputs segment-level copy number, corresponding bin-level estimates for each tool were obtained by intersecting the bins with PURPLE’s segments, then computing the mean PURPLE copy number of all intersecting segments for each bin weighted by the lengths of the segment overlaps with the bin. As PURPLE imputes copy number in some regions instead of estimating them directly, bins in any tool for which more than 75% of their width overlapped these blacklisted regions were removed from consideration. Correlation was computed using Pearson correlation.

## Code availability

The code for the ATAClone is available at: https://github.com/TrigosTeam/ATAClone

## Data availability

The replicate Kidney Cancer datasets were provided to us by 10X Genomics. The 10k Human PBMCs, Multiome v1.0, Chromium Controller dataset was downloaded from the 10X datasets portal at https://www.10xgenomics.com/datasets/.

The scmixology2 lung cancer cell line mixture dataset and its ground-truth labels were downloaded from NCBI GEO under accession number GSE142285. Additional data was also provided to us by the authors.

The prostate cancer matched 10X Genomics single-nuclei multiome and WGS data is available in the European Genome-Phenome Archive under accession number EGAD50000001869. Preprocessing of this data is described in (23).

## Authors’ contributions

LC conceived the development of the tool, developed scientific rationale and statistics of the tool, designed analyses, performed analyses, interpreted results, wrote and edited the manuscripts. AST supervised the work, developed research questions, designed analyses, interpreted results, edited the manuscript. All authors read and approved the final manuscript.

## Acknowledgements

We would like to thank Sirui Weng, the Molecular Genomics Core (RRID:SCR_025695), the Peter Mac Research Computing Facility, the Models of Cancer Translational Research Centre, the Bioinformatics Core (RRID: SCR_025901), and the CASCADE program team (Shahneen Sandhu). We would also like to acknowledge 10X Genomics for sharing the full/raw data for the kidney cancer replicate datasets, and Matt Ritchie and Rheza Ghamsari for sharing the full dataset for scmixology2.

## References

1. Schmid, K.T., Symeonidi, A., Hlushchenko, D., Richter, M.L., Tijhuis, A.E., Foijer, F. and Colome-Tatche, M. (2025) Benchmarking scRNA-seq copy number variation callers. Nat Commun, 16, 8777.

2. Fan, J., Lee, H.O., Lee, S., Ryu, D.E., Lee, S., Xue, C., Kim, S.J., Kim, K., Barkas, N., Park, P.J. et al. (2018) Linking transcriptional and genetic tumor heterogeneity through allele analysis of single-cell RNA-seq data. Genome Res, 28, 1217–1227.

3. Feng, X., Zheng, J., Peng, S., Jiang, A., Ng, K.H., Lyu, C., Jin, Q. and Chen, L. (2026) CNAScope: pan-cancer copy number aberration database with functional annotation and interactive visualization. Nucleic Acids Res, 54, D1364–D1375.

4. Serin Harmanci, A., Harmanci, A.O. and Zhou, X. (2020) CaSpER identifies and visualizes CNV events by integrative analysis of single-cell or bulk RNA-sequencing data. Nat Commun, 11, 89.

5. Patel, A.P., Tirosh, I., Trombetta, J.J., Shalek, A.K., Gillespie, S.M., Wakimoto, H., Cahill, D.P., Nahed, B.V., Curry, W.T., Martuza, R.L. et al. (2014) Single-cell RNA-seq highlights intratumoral heterogeneity in primary glioblastoma. Science, 344, 1396–1401.

6. Su, D., Peters, M., Soltys, V. and Chan, Y.F. (2025) Copy number normalization distinguishes differential signals driven by copy number differences in ATAC-seq and ChIP-seq. BMC Genomics, 26, 306.

7. Cadieux, E.L., Mensah, N.E., Castignani, C., Tanić, M., Wilson, G.A., Dietzen, M., Dhami, P., Vaikkinen, H., Verfaillie, A., Martin, C.C., et al. (2022) Copy number-aware deconvolution of tumor-normal DNA methylation profiles. bioRxiv, 2020.2011.2003.366252.

8. Schneider, M.P., Cullen, A.E., Pangonyte, J., Skelton, J., Major, H., Van Oudenhove, E., Garcia, M.J., Chaves Urbano, B., Piskorz, A.M., Brenton, J.D., et al. (2024) scAbsolute: measuring single-cell ploidy and replication status. Genome Biol, 25, 62.

9. Wang, R., Lin, D.Y. and Jiang, Y. (2020) SCOPE: A Normalization and Copy-Number Estimation Method for Single-Cell DNA Sequencing. Cell Syst, 10, 445–452 e446.

10. Zaccaria, S. and Raphael, B.J. (2021) Characterizing allele- and haplotype-specific copy numbers in single cells with CHISEL. Nat Biotechnol, 39, 207–214.

11. Kolbeinsdottir, S., Zachariadis, V., Sommerauer, C., Lohi, O., Heinaniemi, M. and Enge, M. (2025) Absolute copy number aware CNV calling of sub-megabase segments in ultra-low coverage single-cell DNA sequencing data. Nucleic Acids Res, 53.

12. Ramakrishnan, A., Symeonidi, A., Hanel, P., Schmid, K.T., Richter, M.L., Schubert, M. and Colome-Tatche, M. (2023) epiAneufinder identifies copy number alterations from single-cell ATAC-seq data. Nat Commun, 14, 5846.

13. Nikolic, A., Singhal, D., Ellestad, K., Johnston, M., Shen, Y., Gillmor, A., Morrissy, S., Cairncross, J.G., Jones, S., Lupien, M. et al. (2021) Copy-scAT: Deconvoluting single-cell chromatin accessibility of genetic subclones in cancer. Sci Adv, 7, eabg6045.

14. Wang, X., Jin, Z., Shi, Y. and Xi, R. (2025) Detecting copy-number alterations from single-cell chromatin sequencing data by AtaCNA. Cell Rep Methods, 5, 100939.

15. Takeuchi, F. and Kato, N. (2024) Ploidy inference from single-cell data: application to human and mouse cell atlases. Genetics, 227.

16. Moore, T.W. and Yardımcı, G.G. (2023) Robust CNV detection using single-cell ATAC-seq. bioRxiv, 2023.2010.2004.560975.

17. Gao, R., Bai, S., Henderson, Y.C., Lin, Y., Schalck, A., Yan, Y., Kumar, T., Hu, M., Sei, E., Davis, A. et al. (2021) Delineating copy number and clonal substructure in human tumors from single-cell transcriptomes. Nat Biotechnol, 39, 599–608.

18. Gao, T., Soldatov, R., Sarkar, H., Kurkiewicz, A., Biederstedt, E., Loh, P.R. and Kharchenko, P.V. (2023) Haplotype-aware analysis of somatic copy number variations from single-cell transcriptomes. Nat Biotechnol, 41, 417–426.

19. Buenrostro, J.D., Wu, B., Chang, H.Y. and Greenleaf, W.J. (2015) ATAC-seq: A Method for Assaying Chromatin Accessibility Genome-Wide. Curr Protoc Mol Biol, 109, 21 29 21–21 29 29.

20. Satpathy, A.T., Granja, J.M., Yost, K.E., Qi, Y., Meschi, F., McDermott, G.P., Olsen, B.N., Mumbach, M.R., Pierce, S.E., Corces, M.R., et al. (2019) Massively parallel single-cell chromatin landscapes of human immune cell development and intratumoral T cell exhaustion. Nat Biotechnol, 37, 925–936.

21. Technical Note – Chromium Nuclei Isolation Kit: Data Highlights & Methods Comparison for Single Cell Multiome ATAC + Gene Expression, Document Number CG000552 Rev A, 10X Genomics, (August 16, 2022).

22. Tian, L., Jabbari, J.S., Thijssen, R., Gouil, Q., Amarasinghe, S.L., Voogd, O., Kariyawasam, H., Du, M.R.M., Schuster, J., Wang, C., et al. (2021) Comprehensive characterization of single-cell full-length isoforms in human and mouse with long-read sequencing. Genome Biol, 22, 310.

23. Weng, S., Cain, L., Comben, J., Zhang, Y., Semple, T., Alaei, S., Yoannidis, D., Martelotto, L., Mitchell, C., Pasam, A. et al. (2026) Recurrent intra-tumour heterogeneity is a hallmark of metastatic prostate cancer. Nat Commun.

24. Song, Y., Mei, Z., Zheng, Q., Yuan, Q., Liang, Y., Gao, J., Zhou, L., Wu, S. and Wu, W. (2025) Benchmarking Ploidy Estimation Methods for Bulk and Single-Cell Whole Genome Sequencing. Adv Sci (Weinh), 12, e07839.

25. Schmidt, M.J., Naghdloo, A., Prabakar, R.K., Kamal, M., Cadaneanu, R., Garraway, I.P., Lewis, M., Aparicio, A., Zurita-Saavedra, A., Corn, P. et al. (2025) Polyploid cancer cells reveal signatures of chemotherapy resistance. Oncogene, 44, 439–449.

26. Stuart, T., Srivastava, A., Madad, S., Lareau, C.A. and Satija, R. (2021) Single-cell chromatin state analysis with Signac. Nat Methods, 18, 1333–1341.

27. Lun, A.T., McCarthy, D.J. and Marioni, J.C. (2016) A step-by-step workflow for low-level analysis of single-cell RNA-seq data with Bioconductor. F1000Res, 5, 2122.

28. Butler, A., Hoffman, P., Smibert, P., Papalexi, E. and Satija, R. (2018) Integrating single-cell transcriptomic data across different conditions, technologies, and species. Nat Biotechnol, 36, 411–420.

29. Lin, Y., Ghazanfar, S., Strbenac, D., Wang, A., Patrick, E., Lin, D.M., Speed, T., Yang, J.Y.H. and Yang, P. (2019) Evaluating stably expressed genes in single cells. Gigascience, 8.

30. Meuleman, W., Muratov, A., Rynes, E., Halow, J., Lee, K., Bates, D., Diegel, M., Dunn, D., Neri, F., Teodosiadis, A. et al. (2020) Index and biological spectrum of human DNase I hypersensitive sites. Nature, 584, 244–251.

31. Miao, Z. and Kim, J. (2024) Uniform quantification of single-nucleus ATAC-seq data with Paired-Insertion Counting (PIC) and a model-based insertion rate estimator. Nat Methods, 21, 32–36.

32. Lun, A.T.L., Riesenfeld, S., Andrews, T., Dao, T.P., Gomes, T., participants in the 1st Human Cell Atlas, J. and Marioni, J.C. (2019) EmptyDrops: distinguishing cells from empty droplets in droplet-based single-cell RNA sequencing data. Genome Biol, 20, 63.

33. Muskovic, W. and Powell, J.E. (2021) DropletQC: improved identification of empty droplets and damaged cells in single-cell RNA-seq data. Genome Biol, 22, 329.

34. Orchard, P., Kyono, Y., Hensley, J., Kitzman, J.O. and Parker, S.C.J. (2020) Quantification, Dynamic Visualization, and Validation of Bias in ATAC-Seq Data with ataqv. Cell Syst, 10, 298–306 e294.

35. Halstead, M.M., Kern, C., Saelao, P., Chanthavixay, G., Wang, Y., Delany, M.E., Zhou, H. and Ross, P.J. (2020) Systematic alteration of ATAC-seq for profiling open chromatin in cryopreserved nuclei preparations from livestock tissues. Sci Rep, 10, 5230.

36. Lareau, C.A., Ma, S., Duarte, F.M. and Buenrostro, J.D. (2020) Inference and effects of barcode multiplets in droplet-based single-cell assays. Nat Commun, 11, 866.

37. Ahlmann-Eltze, C. and Huber, W. (2023) Comparison of transformations for single-cell RNA-seq data. Nat Methods, 20, 665–672.

38. irlba: Fast Truncated Singular Value Decomposition and Principal Components Analysis for Large Dense and Sparse Matrices. R package version 2.3.7, https://github.com/bwlewis/irlba.

39. Antonov, M., Csárdi, G., Horvát, S., Müller, K., Nepusz, T., Noom, D., Salmon, M., Traag, V., Foucault Welles, B. and Zanini, F. (2023) igraph enables fast and robust network analysis across programming languages. 10.48550/arXiv.2311.10260.

40. Traag, V.A., Waltman, L. and van Eck, N.J. (2019) From Louvain to Leiden: guaranteeing well-connected communities. Sci Rep, 9, 5233.

41. Traag, V.A., Van Dooren, P. and Nesterov, Y. (2011) Narrow scope for resolution-limit-free community detection. Phys Rev E Stat Nonlin Soft Matter Phys, 84, 016114.

42. Germain, P.L., Lun, A., Garcia Meixide, C., Macnair, W. and Robinson, M.D. (2021) Doublet identification in single-cell sequencing data using scDblFinder. F1000Res, 10, 979.

