## Supplementary Figures for "ATAClone: Cancer Clone Identification and Copy Number Estimation from Single-cell ATAC-seq"

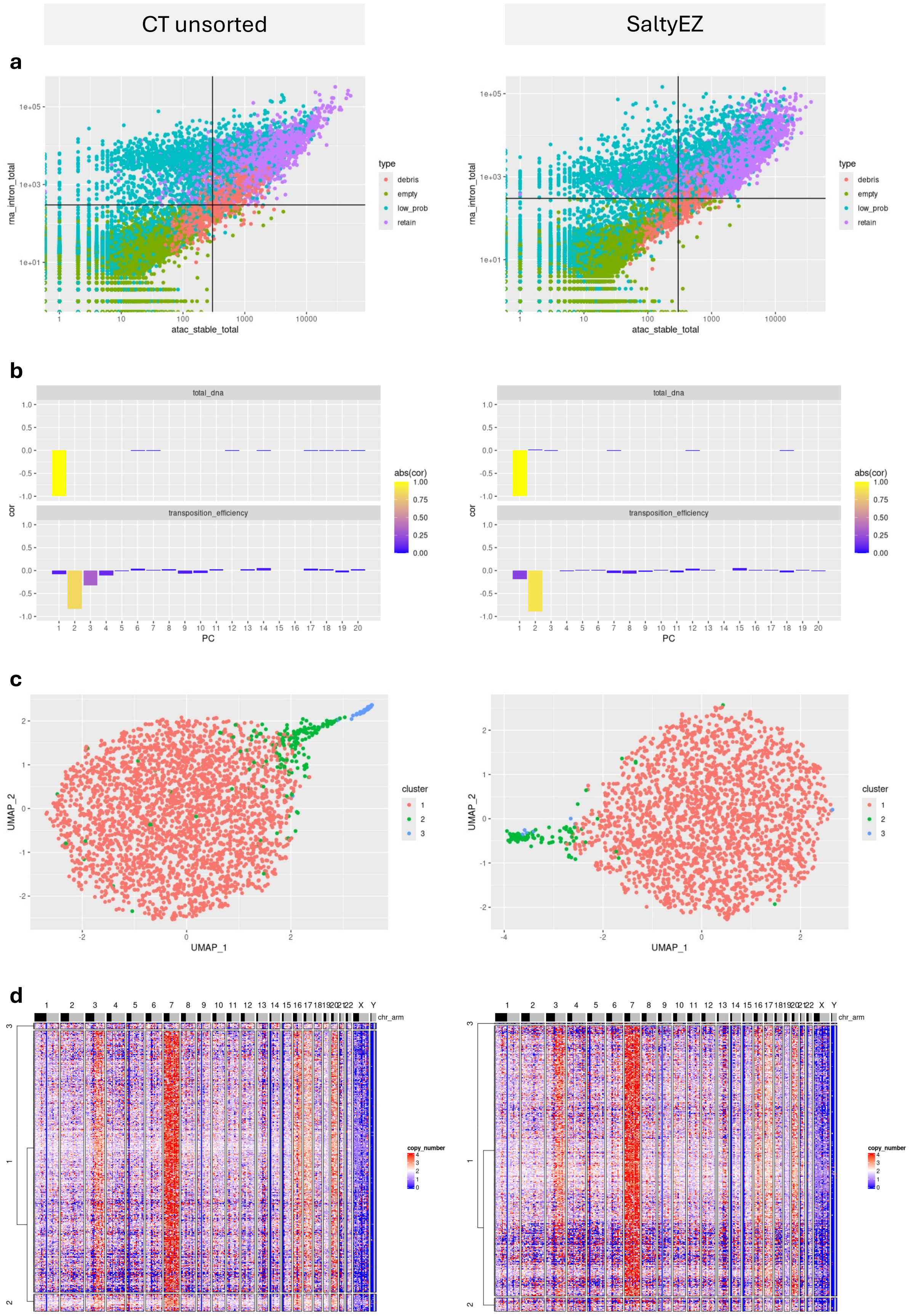


**Supplementary Figure S1. Comparison of ATAClone workflow between 10X Human Kidney Cancer replicate experiments (CT unsorted vs. SaltyEZ)**

Starting from unfiltered data, ATAClone was run to completion for both the CT unsorted (left column) and the SaltyEZ (right column) 10X Human Kidney Cancer replicate experiments. **(a)** Using a standardised set of filtering parameters, ATAClone automatically filters cells for several artefacts. Scatter plots of total ATAC-seq stably-accessible fragments (X axis) vs. total RNA-seq intronic UMIs (Y axis) per cell barcode. Horizontal and vertical lines show the minimum X and Y value to be retained for downstream analysis (**300** for both axes in both experiments). Points are coloured by their classification as artefactual (red = nuclear debris, green = empty droplet, cyan = low probability (under-sequenced) cell barcode) or non-artefactual (purple). Note that low probability barcodes (cyan points) represent the same set of cell barcode sequences in all experiments. Additionally, droplets with a fraction of reads in stably-accessible regions less than 0.05 were also filtered (not shown). **(b)** Bar plots showing Pearson correlation of the first 25 principal components after filtering and normalisation with unwanted technical co-variates. The unwanted technical co-variates shown are total DNA (top) and transposition efficiency (bottom), as measured by the total stably accessible ATAC-seq fragments per barcode and the 'β' statistic (see: Methods), respectively. Bar height represents the correlation with each co-variate (positive or negative) while bar colour represents its absolute value. **(c)** Scatter plot showing UMAP embeddings (X and Y axes) of filtered barcodes and coloured by unsupervised cluster assignment by ATAClone's automated clustering approach. Both UMAP embeddings and clusters are computed based on PCA embeddings of the normalised and filtered data after discarding PCs associated with the technical co-variates shown in **(b)**. **(d)** Heatmap showing 'relative' copy number estimates (i.e. scaling total DNA to be same as reference cells) across consecutive ~10MB bins (columns) for each cell barcode (rows), based on an external set of normal reference cells. Cell barcodes are grouped by the same clusters as in **(c)** with clusters sorted by single-linkage hierarchical clustering on the Manhattan distances of their mean bin values. Clusters of non-tumour cells (Cluster 2 in the CYT unsorted and Clusters 2 and 3 in the SaltyEZ) are distinguishable both as an outgroup in this clustering, and by their relatively constant copy number values. Copy number is clipped to a maximum value of 4 for clearer visualisation.

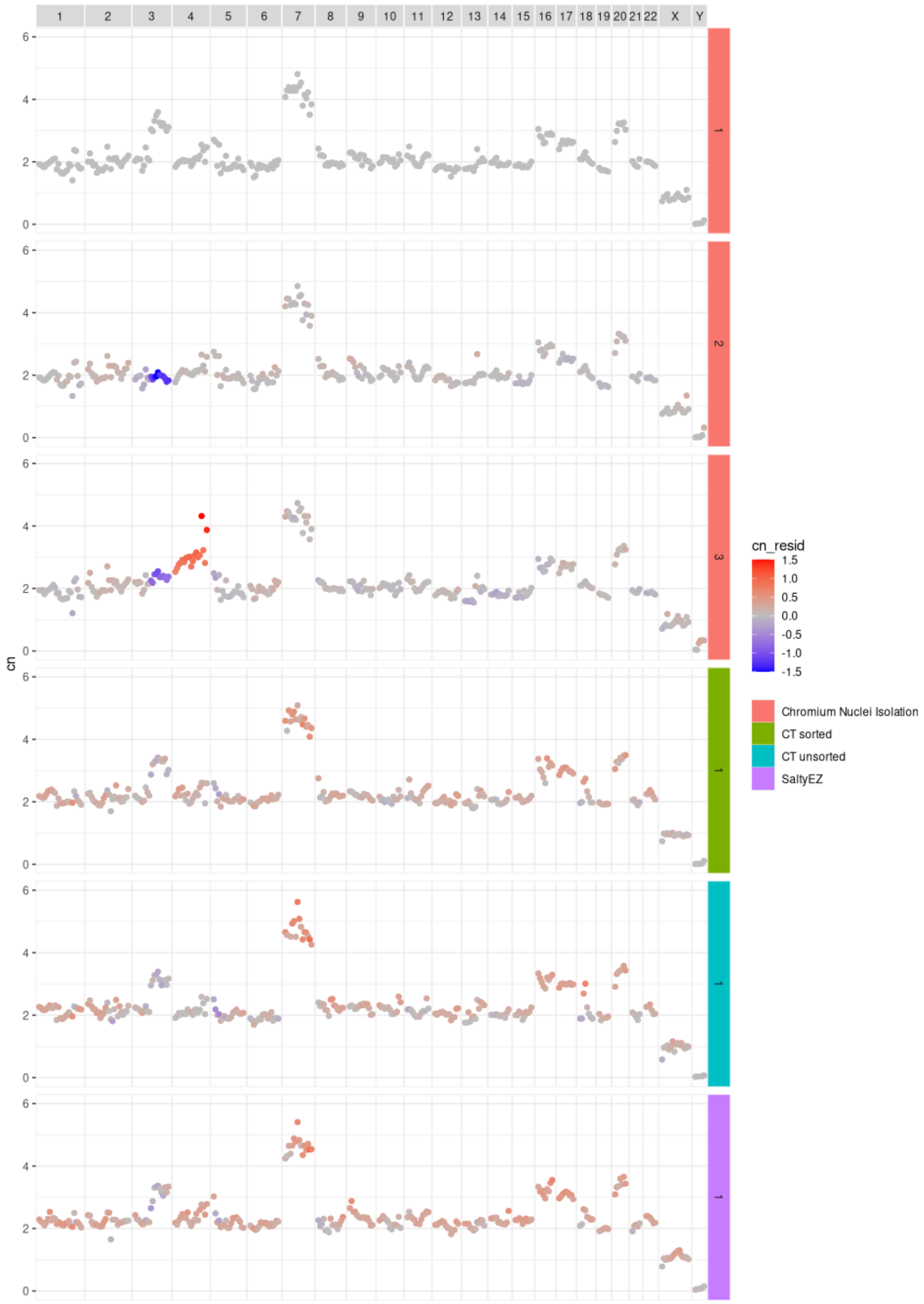


**Supplementary Figure S2. Absolute copy number estimates for all tumour clones across the 10X Human Kidney Cancer replicate experiments.**

Scatter plots show copy number (Y axis) for each consecutive ~10MB bin along the length of each chromosome (X axis) for each clone identified. Label colours identify the experiment the clone belongs to while label numbers correspond to the cluster the clone was assigned to in that experiment (i.e. the same numbers as shown in Figures 2c, 2d, S1c, and S1d). Note that clusters corresponding to either non-tumour cells or doublets were removed. Points are coloured by their difference in estimated copy number from a reference clone (here, Clone 1 in the Chromium Nuclei Isolation experiment is the reference) for the same bin, clipped to ±1.5.

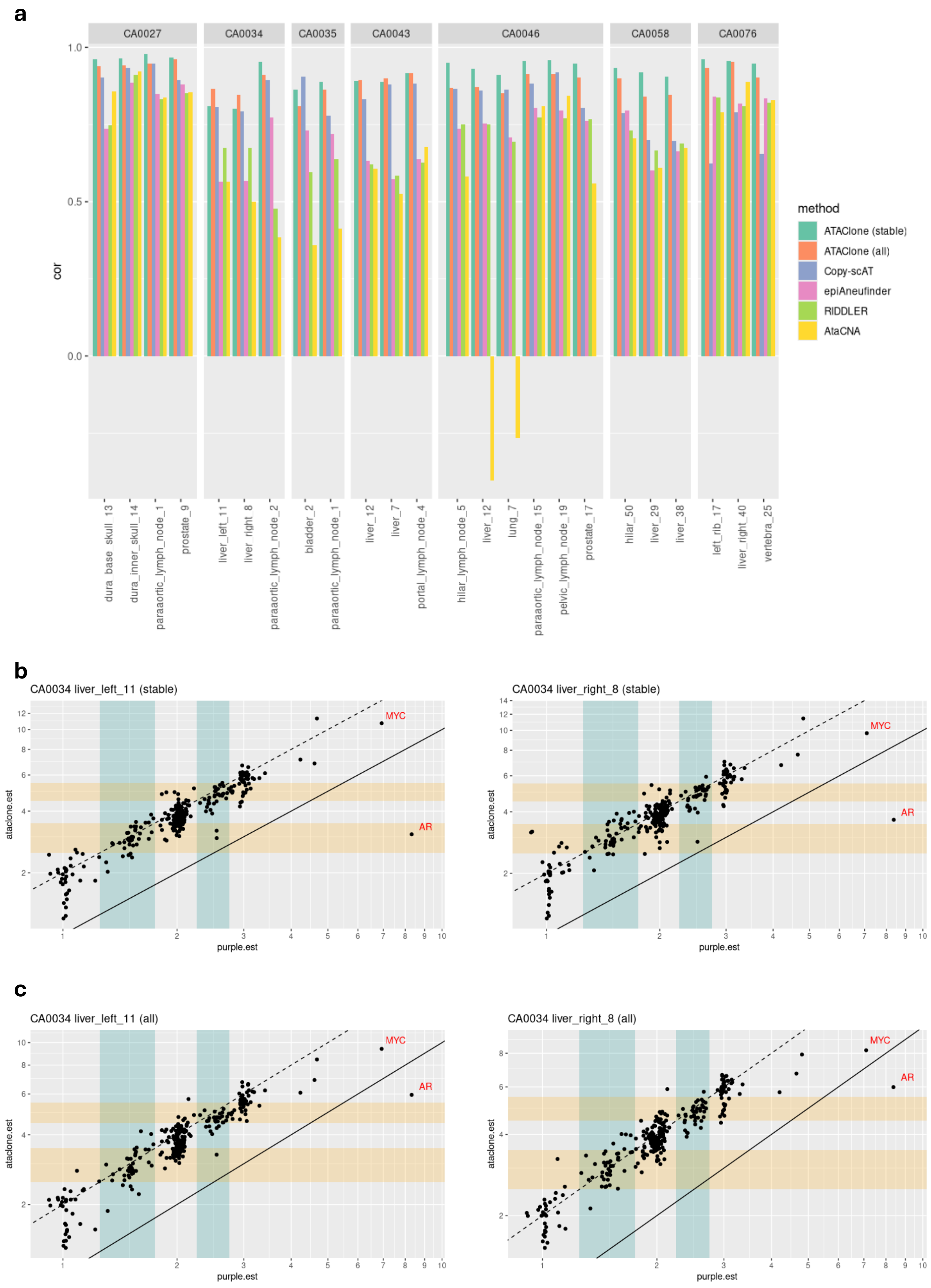


**Supplementary Figure S3: Restriction to stably-accessible regions reduces noise but may introduce some coverage biases**

**(a)** To evaluate the accuracy of ATAClone's copy number estimates, pseudo-bulked copy number estimates from ATAClone, Copy-scat, epiAneudinfer, RIDDLER, and AtaCNA were compared to WGS-derived estimates for matched tissue across 24 metastatic prostate cancer samples using Pearson correlation. Two versions of ATAClone's copy number estimates are shown: estimates based on fragments at stably-accessible regions only (stable) and estimates based on all fragments (all). Correlations are grouped at the facet level by patient identifier (CA- prefix) and by the site within each patient they originate from (x-axis). **(b-e)** While ATAClone generally achieves better correlation with the matched WGS estimates in its "stable" mode, this was not the case for the CA0034 liver_left_11 and CA0034 liver_right_8 samples. These samples in "stable" mode also represent the lowest overall correlation with the matched WGS estimates for ATAClone at 0.81 and 0.80, respectively. To investigate this discrepancy, scatterplots of WGS-derived copy number estimates from PURPLE (x-axis) vs pseudo-bulked single-cell-derived copy number estimates from ATAClone (y-axis) are shown for these samples in "stable" mode **(b)** and "all" mode **(c)**. The bins containing MYC and AR are labelled and correspond to the regions chr8:125150909-135144772 and chrX:61000000-70504090, respectively; the latter displays a notable shift in estimated copy number by ATAClone in "all" mode. Teal vertical bars highlight the inter-integer copy number intervals 1.25-1.75 and 2.25-2.75 (centred on 1.5, and 2.5, respectively) while gold horizontal bars highlight the equivalent doubled intervals 2.5-3.5 and 4.5-5.5 (centred on 3 and 5, respectively). A high density of copy number values in both the teal and gold regions suggests a failure in ploidy estimation by PURPLE which is rectified by ATAClone. Parallel lines correspond to y = x (solid), and y = 2x (dashed). Note that, for visual clarity, all scatterplot axes are displayed on a log scale. However, all correlations displayed in **(a)** were computed in the original linear scale.

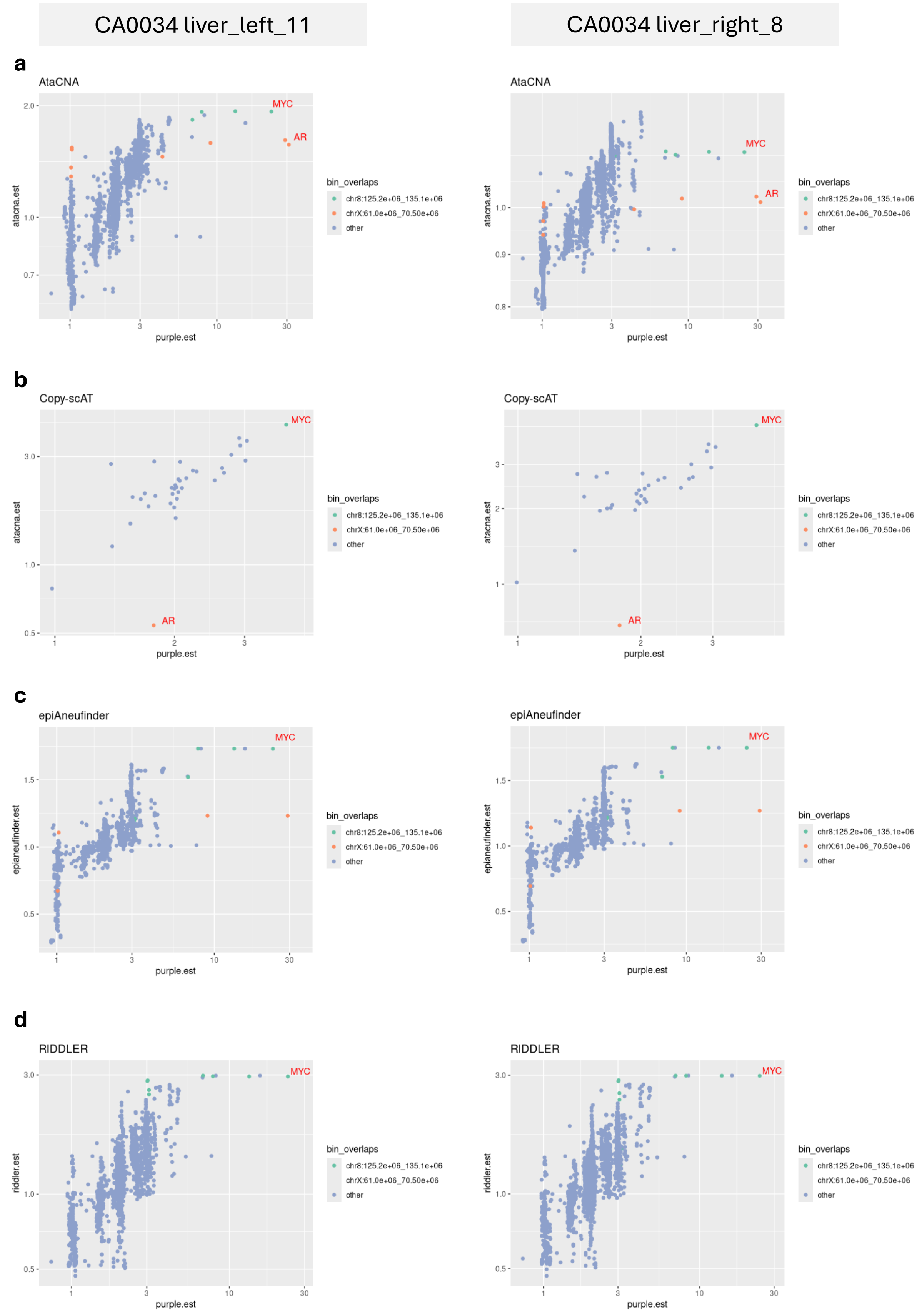


**Supplementary Figure S4. Comparison of oncogene copy number estimation across methods**

**(a-d)** To provide a qualitative comparison of oncogene detection across tools, pseudo-bulked copy number estimates for AtaCNA **(a)**, Copy-scAT **(b)**, epiAneufinder **(c)**, and RIDDLER **(d)** and their corresponding WGS-derived estimates are also shown for the CA0034 liver_left_11 and CA0034 liver_right_8 samples (for the corresponding ATAClone results, see Supplementary Figure S3). Bins intersecting the regions chr8:125150909-135144772 and chrX:61000000-70504090 (i.e., the corresponding ATAClone bins labelled in Supplementary Figure S3) are highlighted while the bins containing the MYC and AR genes themselves are labelled by text, when present. A bin containing AR is absent from epiAneufinder due to its default filtering behaviour and from RIDDLER due to filtering chrX by default; Copy-scAT bins correspond to full chromosome arms. Note that, for visual clarity, all scatterplot axes are displayed on a log scale except for the y axis of the epiAneufinder plots which are linear.
